# Protein binding and methylation on looping chromatin accurately predict distal regulatory interactions

**DOI:** 10.1101/022293

**Authors:** Sean Whalen, Rebecca M. Truty, Katherine S. Pollard

## Abstract

Identifying the gene targets of distal regulatory sequences is a challenging problem with the potential to illuminate the causal underpinnings of complex diseases. However, current experimental methods to map enhancer-promoter interactions genome-wide are limited by their cost and complexity. We present *TargetFinder*, a computational method that reconstructs a cell’s three-dimensional regulatory landscape from two-dimensional genomic features. *TargetFinder* achieves outstanding predictive accuracy across diverse cell lines with a false discovery rate up to fifteen times smaller than common heuristics, and reveals that distal regulatory interactions are characterized by distinct signatures of protein interactions and epigenetic marks on the DNA loop between an active enhancer and targeted promoter. Much of this signature is shared across cell types, shedding light on the role of chromatin organization in gene regulation and establishing *TargetFinder* as a method to accurately map long-range regulatory interactions using a small number of easily acquired datasets.

---

Genotyping, exome sequencing, and whole-genome sequencing have linked thousands of non-coding variants to traits in humans and other eukaryotes [1–6] via genome-wide association studies, family studies, and other approaches. Non-coding variants are more likely to cause common disease than are non-synonymous coding variants [7], and they can account for the vast majority of heritability [8]. Yet few non-coding mutations have been functionally characterized or mechanistically linked to human phenotypes [7, 9]. Comparative [10] and functional [11–13] genomics, coupled with bioinformatics, are generating annotations of regulatory elements in many organisms and cell types [14], as well as tools for exploring or predicting the impact of mutations in regulatory DNA [15–18]. Massively parallel reporter assays [19] provide a way to test and refine some of these predictions. However, this new information will only improve our understanding of disease and other phenotypes if we can accurately link functional non-coding elements to the genes, pathways, and cellular processes they regulate. This is a difficult problem because promoters and their regulatory elements can be separated by thousands—even greater than one million [20]—base pairs (bp), and the closest promoter is usually not the true target in humans [21] (though this varies by species [22]). Incorrectly mapping regulatory variants to genes prevents meaningful downstream studies. Our goal is to address this problem by developing a computational method to accurately predict distal regulatory interactions with relatively easy-to-collect data.

A computational method is desirable for two reasons. First, experimental mapping of chromatin interactions at the resolution of individual promoters and regulatory elements on a genome-wide scale in many cell types and developmental stages is prohibitively expensive and technically challenging. Until recently, very few validated distal regulatory interactions were known. Hence, previous studies defined interactions indirectly via genomic proximity coupled with genetic associations (e.g., eQTLs [23]), gene expression [14, 24–26], or promoter chromatin state [27, 28]. High-throughput methods for assaying chromatin interactions now exist, including paired-end tag sequencing (ChIA-PET) [29] and extensions of the chromosome conformation capture (3C) assay [30] (5C, Hi-C) [31, 32]. Recent improvements to the resolution of genome-wide Hi-C give an unprecedented look at chromatin structure [33, 34], including studies that utilize sequence capture to enrich for interactions with annotated promoters [35]. But Hi-C at the resolution of individual regulatory elements is still prohibitively expensive for most labs. We therefore saw the first high-resolution Hi-C experiments as an opportunity to test the hypothesis that the spatial proximity of promoters with distal regulatory elements can be inferred from data that is routinely measured genome-wide, including DNA sequences, epigenetic marks, and/or protein-DNA binding events. An accurate and generalizable model would enable high-resolution *in silico* Hi-C for many cell types using data that already exists or can be collected rapidly. A second motivation to computationally model regulatory interactions using functional genomics data is to learn relationships between DNA sequences, structural proteins, transcription factors and modified histones in the context of chromatin looping and gene activation. Learning which combinations of features best predict looping might reveal novel protein functions and molecular mechanisms of distal gene regulation.

We implemented an algorithm called *TargetFinder* that integrates hundreds of functional genomics and sequence datasets to identify the minimal subset necessary for accurately predicting enhancer-promoter interactions across the genome. We focused on enhancers due to their large impact on gene regulation [36] and our ability to predict their locations genome-wide, though our approach could easily be extended to other classes of regulatory elements. Applying *TargetFinder* to three human ENCODE cell lines [11] with high resolution Hi-C data [33], we discovered that enhancer-promoter interactions can be predicted with extremely high accuracy. Interestingly, these analyses showed that functional genomics data marking the window *between* the enhancer and promoter are more useful for identifying true interactions than are proximal marks at the enhancer and promoter. Exploration of this phenomenon revealed specific proteins and chemical modifications on the chromatin loop that bring an enhancer in contact with its target promoter and not with nearby repressed or active but non-targeted promoters. Thus, *TargetFinder* provides a framework for accurately assaying three-dimensional genomic interactions, as well as techniques for mining massive collections of experimental data to shed new light on the mechanisms of distal gene regulation.

## Results

### Ensemble learning of regulatory interactions from genomic data

The core component of *TargetFinder* is a machine learning pipeline that builds and evaluates ensemble models of distal regulatory interactions from genomic features such as epigenetic marks, protein binding events, gene annotations, and evolutionary signatures (Figure 1). Ensemble learning methods have excellent performance, account for non-linear interactions between features, and estimate the predictive importance of each feature. We applied multiple ensemble methods, including random forests and gradient boosted trees, to ensure our conclusions are robust. The inputs to *TargetFinder* are pairs of enhancers and promoters, annotated as interacting or not in a given cell type, and features (i.e., summaries of genomic datasets) associated with each pair. The outputs are a model for predicting interactions in new enhancer-promoter pairs, assessments of model performance on held-out data, and measurements of how predictive each feature is alone and in combination with other features. The predictive contribution of different genomic regions is explored by separately quantifying the importance of features marking enhancers, promoters, and the genomic window between them, and we also examine the importance of each feature alone versus in combination with other features. We specifically implemented *TargetFinder* to address the challenging problem of distinguishing validated enhancer-promoter interactions from all non-interacting pairs of transcribed gene promoters and active distal enhancers (> 10 kilobases (Kb) from the transcription start site (TSS)) within any 2 megabase (Mb) locus. The method is easily extended to include other types of regulatory elements, such as inactive promoters and enhancers, but we found that including inactive elements resulted in less informative models.

**Figure 1:**
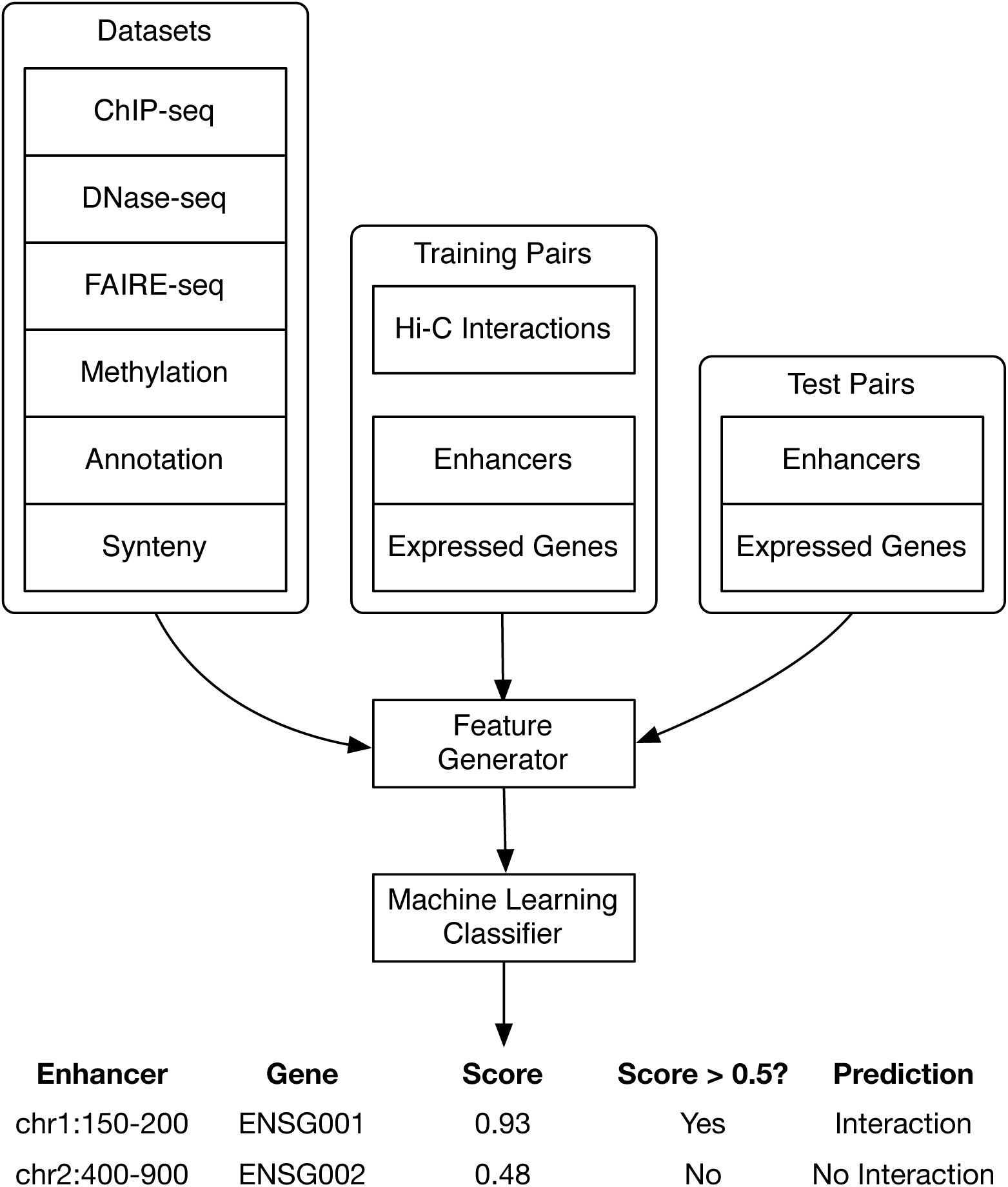
The *TargetFinder* pipeline. Features are generated from hundreds of diverse datasets for pairs of enhancers and promoters of expressed genes found to have significant Hi-C interactions (positives), as well as random pairs of enhancers and promoters without significant interactions (negatives). These labeled samples are used to train an ensemble classifier that is used to examine the predictive importance of each feature or predict whether new enhancer-promoter pairs interact. Classifier predictions are probabilities, and a decision threshold (commonly 0.5 but may be adjusted) converts these to positive or negative prediction labels. Though any classifier can be used, we selected two popular ensemble methods (random forests and gradient boosted trees; Methods) for their predictive accuracy and interpretability. This figure excludes the selection of minimal predictor sets for simplicity.

### TargetFinder predicts enhancer-promoter pairs with high accuracy

We first identified active promoters and enhancers in three ENCODE cell lines that have rich functional genomics data as well as interaction data produced by Rao et al. [33]. These included K562 (mesoderm lineage cells from a leukemia patient), GM12878 (lymphoblastoid cells), and HeLa-S3 (ectoderm lineage cells from a cervical cancer patient). Datasets included measures of open chromatin, DNA methylation, and chromatin immunoprecipitation followed by sequencing (ChIP-seq) for transcription factors (TFs), architectural proteins, and modified histones (Supplemental Table S2). We also included features representing conserved synteny (occurring nearby across species) and co-annotation of the target gene and TFs with motifs in the enhancer (Methods). Using high resolution genome-wide measurements of chromatin interactions [33] in these lines, we annotated interacting and non-interacting enhancer-promoter pairs and generated their corresponding features. For each individual cell line as well as their combination, we repeatedly built models using a random subset of the data and then quantified predictive accuracy on the held-out data using various metrics (Methods), including a balance of precision and recall (power) called *F*_max_. Due to the large number of non-interacting pairs, precision and recall are less biased than the commonly reported area under the receiver operating characteristic curve (AUC) that de-emphasizes false positives.

*TargetFinder* performed well on held-out data in all cell lines, achieving *F*_max_ between 83-88% corresponding to 5-12% false discovery rate (FDR) or approximately 87% power at a 10% false positive rate (Table 1). Performance was similar across lines, with *TargetFinder* performing slightly worse in GM12878 compared to K562 and HeLa-S3. This variability is due in part to differences in the number of training samples as well as the quality and quantity of functional genomics data. Performance was nearly identical using random forests and gradient boosting, and both performed significantly better than non-ensemble methods. Since gradient boosting provides more options for estimating feature importance, subsequent results are from this algorithm. Interestingly, we found *TargetFinder* has very high precision and recall largely independent of enhancer-promoter interaction distances in the range of 10 Kb to 2 Mb (Supplementary Figures S1 and S2). By comparison, all commonly used bioinformatics methods have much higher FDR. For example, using the closest active gene has an estimated FDR of 53-77% [21, 37, 38].

**Table 1:** TargetFinder performance on held out data. Metrics include precision, recall, the maximum harmonic mean of precision and recall over all scoring thresholds (*F*_max_), Matthews correlation coefficient (*ϕ*), area under the ROC curve, and power (true positive rate) at a 10% false positive rate. Ensemble methods (random forests and gradient boosting) have similarly high precision and recall compared to linear models due to their ability to capture non-linear feature interactions. The gap between AUC and precision/recall measures demonstrates how the former is biased due to de-emphasis of false positives. Metrics were computed on predictions generated for the test split of each cross-validation fold and then averaged. Precision and recall were computed using the *F*_max_ threshold. Baseline performance was estimated using random training labels.

| | Model | AUC | TPR | Precision | Recall | $F_{\max}$ | MCC |
| --- | --- | --- | --- | --- | --- | --- | --- |
| Cell Line |  |  |  |  |  |  |  |
| K562 | Baseline | 0.50 | 0.00 | 0.17 | 1.00 | 0.29 | 0.01 |
| K562 | Logistic Regression | 0.79 | 0.48 | 0.48 | 0.56 | 0.51 | 0.42 |
| K562 | Gradient Boosting | 0.95 | 0.87 | 0.88 | 0.80 | 0.84 | 0.82 |
| K562 | Random Forest | 0.95 | 0.87 | 0.92 | 0.80 | 0.85 | 0.83 |
| GM12878 | Baseline | 0.50 | 0.00 | 0.17 | 1.00 | 0.29 | 0.00 |
| GM12878 | Logistic Regression | 0.78 | 0.43 | 0.42 | 0.60 | 0.49 | 0.39 |
| GM12878 | Gradient Boosting | 0.94 | 0.85 | 0.87 | 0.76 | 0.81 | 0.79 |
| GM12878 | Random Forest | 0.94 | 0.85 | 0.90 | 0.76 | 0.83 | 0.79 |
| HeLa-S3 | Baseline | 0.49 | 0.00 | 0.17 | 1.00 | 0.29 | -0.02 |
| HeLa-S3 | Logistic Regression | 0.81 | 0.51 | 0.50 | 0.58 | 0.53 | 0.46 |
| HeLa-S3 | Gradient Boosting | 0.96 | 0.89 | 0.93 | 0.81 | 0.87 | 0.84 |
| HeLa-S3 | Random Forest | 0.96 | 0.89 | 0.95 | 0.83 | 0.88 | 0.85 |
| Combined | Baseline | 0.50 | 0.00 | 0.17 | 1.00 | 0.29 | -0.00 |
| Combined | Logistic Regression | 0.76 | 0.41 | 0.39 | 0.59 | 0.47 | 0.36 |
| Combined | Gradient Boosting | 0.94 | 0.86 | 0.86 | 0.77 | 0.81 | 0.80 |
| Combined | Random Forest | 0.95 | 0.87 | 0.91 | 0.77 | 0.83 | 0.80 |

### Variable importance highlights key datasets for predicting interactions

*TargetFinder* quantifies the importance of each feature and annotates whether it is associated with interacting or non-interacting enhancer-promoter pairs. This enabled us to deeply explore the genomic data associated with chromatin loops and revealed several interesting patterns. First, among hundreds of diverse features, the most predictive were functional genomics experiments including specific ChIP-seq, DNase-seq, and DNA methylation experiments (Figure 2). We observed that features differed in importance across cell lines for many reasons, including real functional differences (e.g., different co-factors), lack of expression (e.g., tissue-specific TFs), data quality, and the experimental design of ENCODE (including data processing methods and availability of replicates). Combining cell lines, the most predictive features were DNA methylation and binding of structural proteins and histones. Proteins related to activation (in particular, activator protein 1 (AP-1) complex [39]) and repression (in particular, polycomb repressive complex 2 (PRC2) [40]) also boost performance. These results demonstrate the rich information about chromatin looping that is present in datasets that are easier and less costly to collect than high-resolution Hi-C.

**Figure 2:**
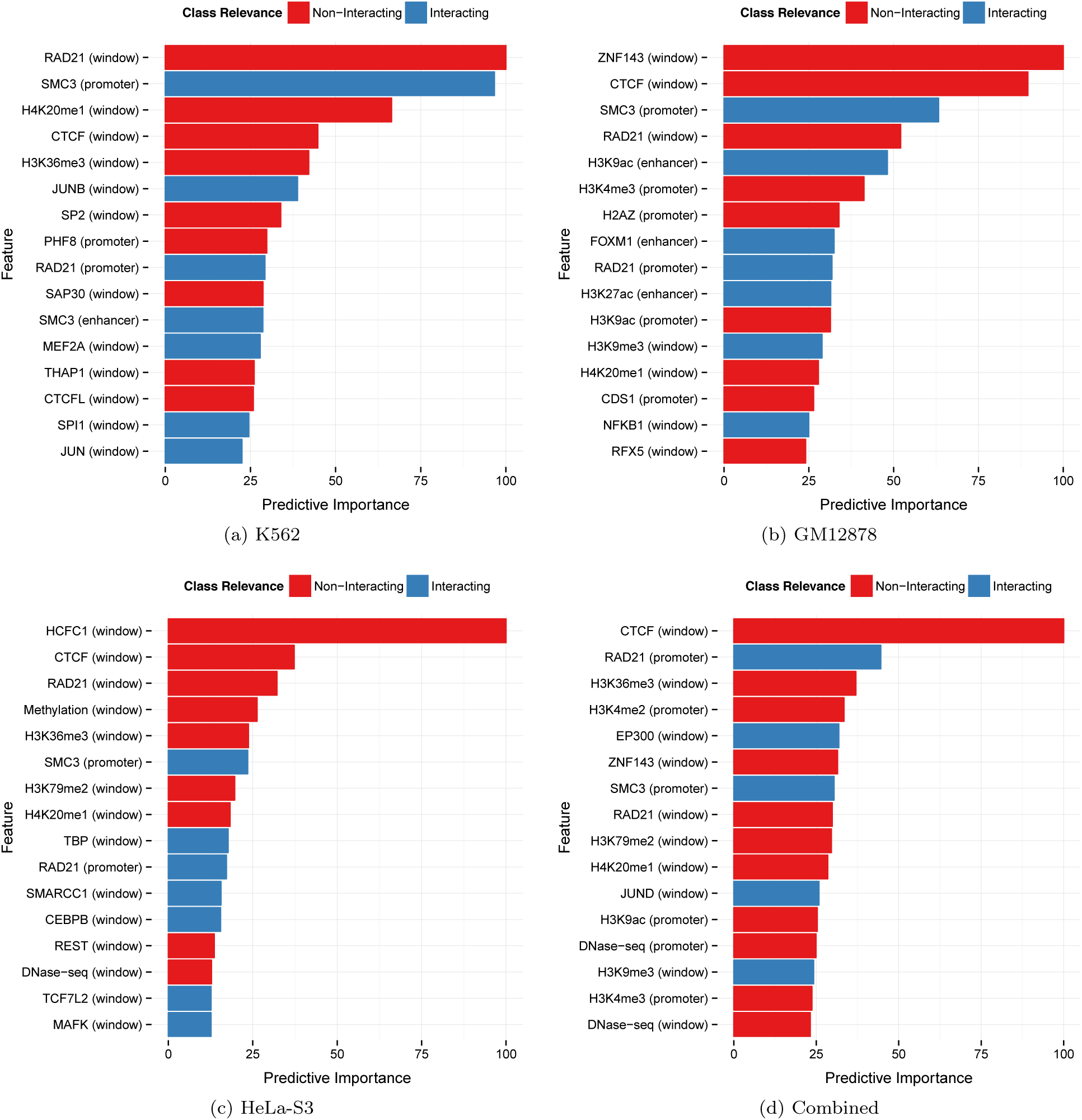
Top 16 predictive features per cell line. Color indicates relevance to interacting (blue) or non-interacting (red) enhancer-promoter pairs, estimated by a gradient boosting classifier. Features at the region between the enhancer and promoter (the *window*) are more prevalent than those directly at the enhancer and promoter. The same feature (e.g., CTCF, cohesin complex) may be relevant to either interacting or non-interacting pairs depending on whether it binds in the window versus at enhancers or promoters.

The most surprising discovery is that, contrary to our prior belief that *TargetFinder* would mostly utilize proximal marks at the enhancer and promoter, the most informative features were instead protein binding events and epigenetic modifications in the genomic window between an enhancer and its target (Figure 3). This is true despite the fact that average signal (e.g., ChIP-seq peak density) was higher at enhancers and promoters for most features.

**Figure 3:**
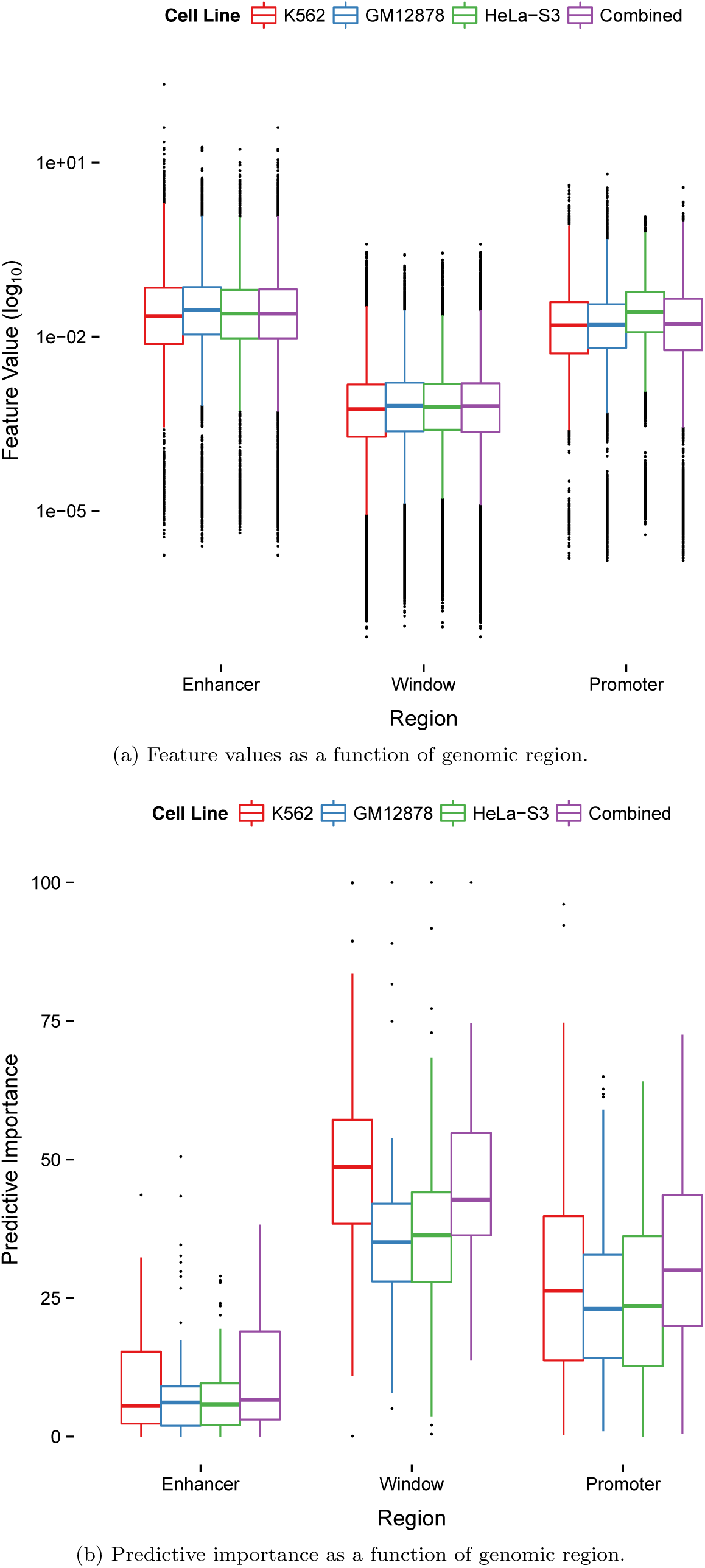
Feature values and predictive performance for enhancer, promoter, and window regions. Despite having the lowest feature values, the predictive importance of the window dominates that of enhancer and promoter regions.

Window features are highly informative for several reasons. Some window features are directly involved in chromatin looping including CTCF, the cohesin complex (SMC3/RAD21), and zinc finger protein ZNF143. The latter interacts with CTCF to provide sequence specificity for chromatin interactions [41], and also interacts with lineage-specific TFs that ZNF143 binds at interacting promoters (e.g., HCFC1 in HeLa-S3 [42]). Other window features impact the likelihood that additional promoters in the locus are the true targets of an enhancer. For example, RNA polymerase II (Pol II) at a promoter is not informative alone, because it can indicate either active transcription or a gene that is poised for rapid activation. In the paused state, Pol II blocks premature activation by acting as an insulator [43]. *TargetFinder* learned that non-targets can be easily distinguished by a lack of activators or coactivators [44] as well as histone marks, such as H3K36me3 and H3K79me2, that are associated with elongation. When these features occur in the window between an enhancer and a promoter, they indicate that an intervening promoter may be the true target. On the other hand, the presence of heterochromatin, PRC2 silencing [40], and various insulators (including existing looping interactions [45] marked by cohesin) in the window suggest that intervening genes are unavailable for binding. Window-associated marks may also be proxies for relevant but un-assayed histone modifications [46]. Thus, functional genomics data in the window between an enhancer and promoter carries rich information about the chromatin conformation of the locus that *TargetFinder* utilizes to predict if the enhancer and promoter physically interact.

Finally, we observed that many of the most important features mark non-interacting enhancer-promoter pairs. In other words, the absence of one feature may be more predictive of looping chromatin than the presence of a different feature. One example is the absence of activators in the window, which is more informative than the presence of Pol II, which might be paused. We also found that the association of a feature with interacting versus non-interacting pairs may be different at promoters, enhancers, and windows between these (Figure 2). For instance, SMC3 at promoters and enhancers is positively associated with interactions, while SMC3 binding in the window region is associated with non-interacting enhancer-promoter pairs because it increases the likelihood that an intervening promoter is the true target. The same holds for histones associated with activation, elongation, and repression.

### TargetFinder identifies complex interactions between DNA-binding proteins and epigenetic marks

There are many top ranked features with similar predictive power. This is due in part to strong correlations between feature values, for example, due to multi-protein complexes or groups of related histone modifications with similar binding patterns across enhancer-promoter pairs (Figure 4). Correlated blocks of top ranked features fall into several broad categories including architectural proteins (CTCF, RAD21, SMC3, ZNF143), DNA methylation, and several types of histone modifications related to elongation (H3K36me3, H3K79me2), heterochromatin (H4K20me1) and activation (H2AZ, H3K4me1/2/3, H3K9ac, H3K27ac). These clusters of features often divide into sub-blocks such that one is associated with interactions and the other with non-interactions, suggesting different roles in chromatin organization despite correlated genomic distributions.

**Figure 4:**
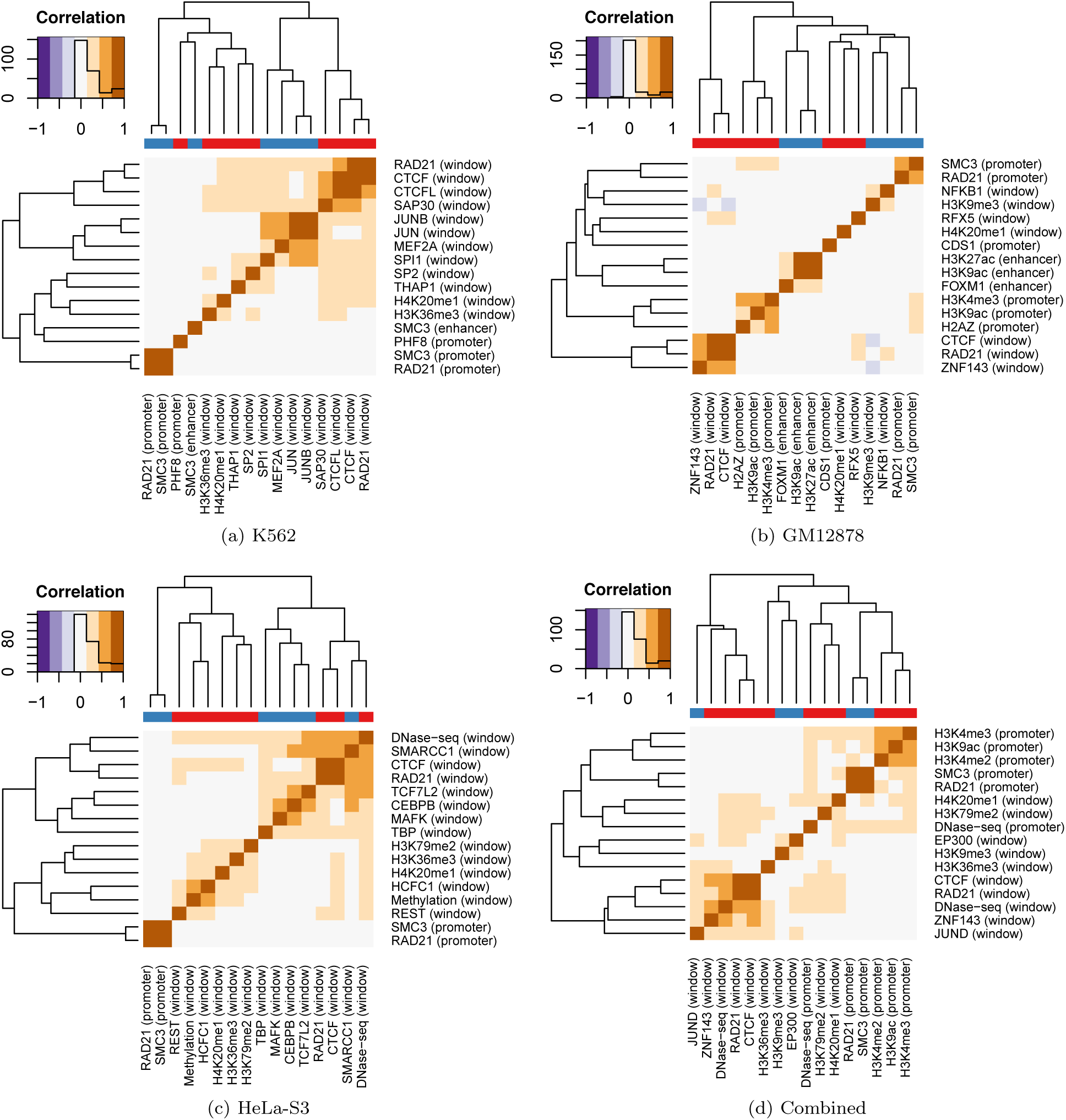
Predictive features co-occur on the genome. Clustered correlation heatmap of the 16 most predictive features per cell line. The color bar along the top of the heatmap indicates feature relevance to interacting (blue) or non-interacting (red) enhancer-promoter pairs. Blocks of correlated predictors along the diagonal often indicate proteins found in the same complex, or histones interacting with a chemical modifier such as an acetyltransferase or deacetylase.

One consequence of correlated features is that proteins performing multiple distinct functions tend not to be highly predictive on their own. Instead, *TargetFinder* preferentially uses their co-factors that specify function. For example, the histone acetyltransferase EP300 is not always a top ranked feature, despite being strongly associated with active enhancers due to its ability to acetylate H3K27 [47]. The importance of EP300 is reduced in some cell lines because it is highly correlated with more predictive co-factors. One such co-factor is C/EBP*β* that has been shown to phosphorylate and modulate the activity of EP300, as well as translocate it to specific gene regions [48]. The predictive importance of ZNF143 is also sometimes reduced due to its correlation with CTCF and the cohesin complex. Considered independent of these other features, however, ZNF143 is among the top 10 predictors for each cell line. Similarly, JUN typically ranks below its co-factors, but has higher importance when *TargetFinder* is trained using all cell lines. This is because the combined model can only utilize features present in all cell lines, and the more predictive co-factors of JUN are not uniformly assayed by ENCODE.

To further explore correlations between co-localizing proteins, we systematically compared how well each feature predicts enhancer-promoter interactions in isolation against its performance in combination with other features (Supplementary Tables S3 to S6). In K562, large changes in predictive rank included SPI1 (PU.1) that has been linked to chromatin looping [49, 50], NCOR1 which represses transcription by inhibiting Pol II elongation [51], TBP that has been linked with long range interactions [52] and whose TAF3 subunit is recruited by CTCF to distal promoters [53], and SRF which regulates FOS [54] and interacts with C/EBP*β* [55]. Large HeLa-S3 rank changes included TBP (see previous), TCF7L2 that is known to form DNA loops by bending [56], SMARCC1 (BAF155) that is part of the SWI/SNF complex implicated in long range looping [57], MAFK that is a subunit of NF-E2 linked to long range interactions and *β*-globin activation [58], and STAT1 linked to chromatin remodeling of the MHC locus [59]. GM12878 had large rank changes for FOXM1 and EBF1 having known roles in B cell fate [60], as well as NFATC1 involved in enhancer-promoter communication [61]. Other large rank changes commonly included activating histone marks such as H2AZ and H3K9ac that may help distinguish active enhancers and promoters, including non-targets within window regions that cannot be discriminated solely by activation marks at their promoters. The elevated importance of H2AZ might also be explained by the link between H2A ubiquitination and polycomb silencing [62]. Thus, changes in predictive rank recapitulate known protein interactions and can identify under-appreciated or novel biological interactions. Lineage-specific proteins without large rank changes also have strong evidence of their relevance to enhancer-promoter interactions, such as RFX5 that can interact with SMC3 to tether looping enhancers to their targets [63].

Next, we combined the correlations shown in Figure 4 with protein-protein interaction data to derive a network of highly predictive features for each cell line that sheds light on the biological interplay between different clusters of regulatory proteins (Figure 5). This provided further support for the observation that histone modifications shared between diverse cell lines are highly predictive when combined with tissue-specific TFs. For example, *TargetFinder* learned that H4K20me1 interacts with H3K9me1/2/3, and this combination is known to mark different types of silent chromatin [64]. It also identified the known interaction between PHF8 and H4K20, which is demethylated by PHF8 during cell cycle progression [65]. These observations are consistent with the finding that histone modifications alone cannot drive transcription, even when Pol II is successfully recruited [66]. Instead, our results underscore that the interaction of histone-modifying complexes such as deacetylases, acetyltransferases, demethylases, and methyltransferases are essential to predicting and understanding the mechanisms of distal gene regulation [67]. A powerful advantage of the tree-based ensemble classifiers used in *TargetFinder* is their ability to detect and utilize these complex, non-linear interactions between features. No single dataset or simple rule captures the genomic signature of all interactions, but *TargetFinder* can learn the more intricate rules needed for distinguishing the true target of an active enhancer (Figure 6, Supplemental Figure S3).

**Figure 5:**
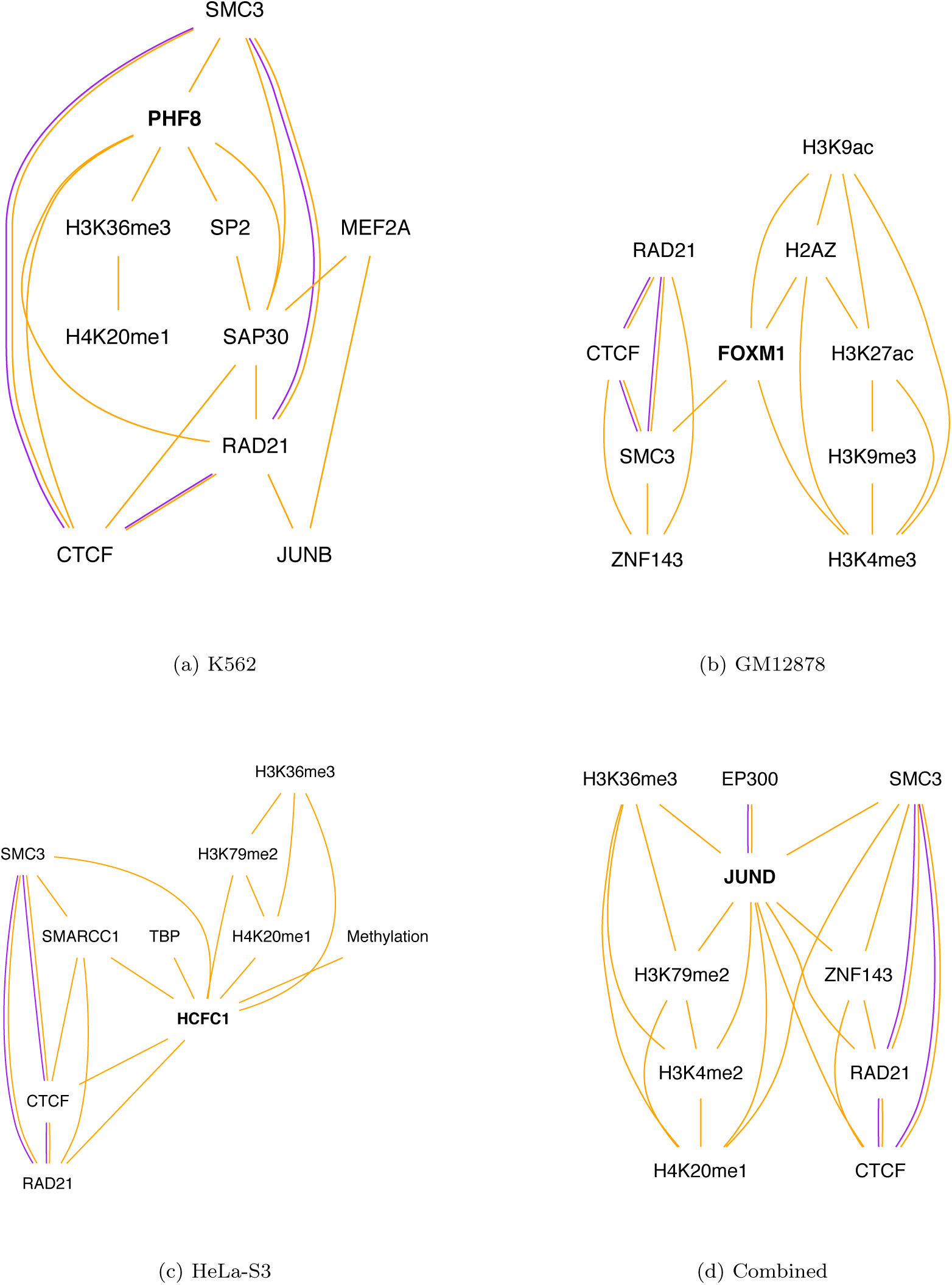
Predictive features physically interact in protein complexes. Interaction network for the top 10 most predictive features per cell line. Purple edges indicate known protein interactions according to BioGrid (Methods). Other features co-occur with these known protein interactions (orange edges indicating moderate to high correlation of peak locations along the genome) and may form complexes with them. The node that is part of the most shortest paths between any two nodes (the most *central* node) is shown in bold and is often a lineage-specific TF.

**Figure 6:**
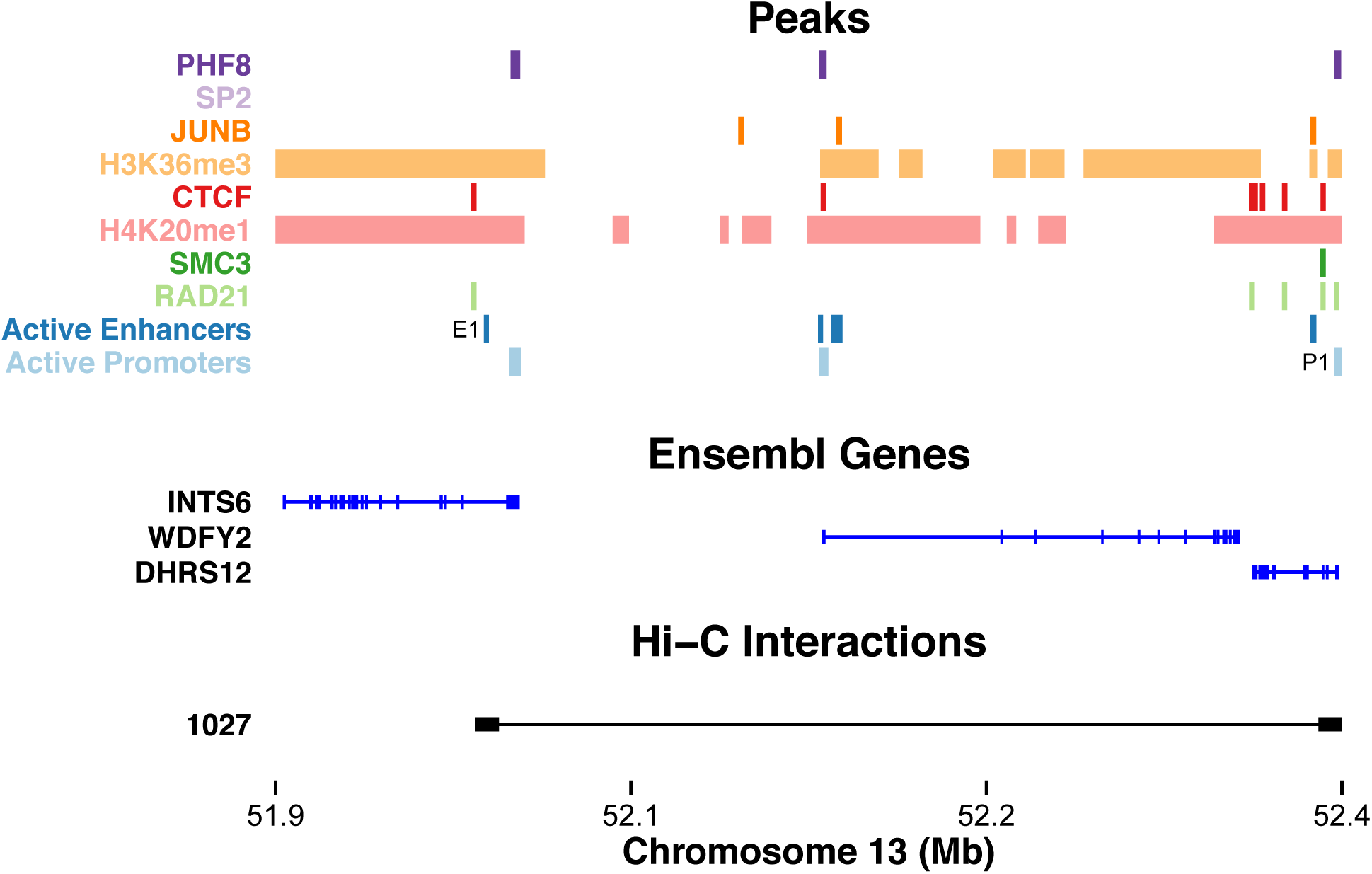
Predicting a chromatin loop that skips over two active promoters in K562 cells. Browser-like tracks display the top 8 predictive datasets (significant peak values), ChromHMM enhancers and promoters, Ensembl genes (introns = thin lines, exons = squares), and Hi-C interactions (left and right fragments connected by a line). An enhancer (E1) interacts with a promoter of DHRS12 (P1) nearly 400 kilo bases away and not with the intervening active promoters of INTS6 and WDFY2. *TargetFinder* integrates multiple datasets (including ones beyond the top 8 displayed) to learn the architecture of this locus. It appears likely that WDFY2 is not targeted due to the presence of the insulator CTCF without interaction-associated cohesin complex marks (SMC3/RAD21), while DHRS12 is marked by RAD21 at the promoter and SMC3/RAD21 nearby. Interestingly, P1 has marks for both H4K20me1 (repression) and H3K36me3 (elongation leading to activation), and the highly predictive mark SP2 is not present in this particular region. These characteristics underscore the need for a machine-learning approach to integrate complex genomic signatures for accurate target prediction.

### TargetFinder generalizes across cell types

We developed and validated *TargetFinder* on the data-rich cell lines provided by ENCODE with an eye towards its application to cell lines or tissues with more limited data, including those without validated enhancer-promoter interactions necessary for training a new model. To explore how well *TargetFinder* generalizes to other cell lines, we used models trained on K562, GM12878, and HeLa-S3 to make predictions on each of the other lines using only the top 16 features from the training cell line that were also available in the test cell line. We also made predictions on the HUVEC line (human umbilical epithelial cells), which has only 11 available features (Supplemental Table S2). Despite the significant differences between cell lines noted above, we found that training *TargetFinder* on one cell line and predicting on another identified a large portion of validated enhancer-promoter interactions (Supplementary Table S1). Estimates of *F*_max_ ranged between 38 and 46%, including on HUVEC, which indicates reasonable precision and recall (e.g., an *F*_max_ of 44% corresponded to 36% precision and 55% recall). Such performance may be sufficient to discover novel enhancer targets without the considerable expense or labor of high-resolution Hi-C experiments.

Since functional genomics data is both highly predictive and costly to obtain, we also evaluated the cross-validation performance of *TargetFinder* using a small number of features. For all cell lines, *TargetFinder* needed ∼16 features to achieve nearly optimal performance (*F*_max_ 0.76-0.81, Figure 7), and comparable performance was achieved with just 8 features (*F*_max_ 0.70-0.77). Thus, regulatory interactions in a new cell type could be predicted by generating less than ten ChIP-seq datasets and using a *TargetFinder* trained on cell types where validated interactions have already been obtained.

**Figure 7:**
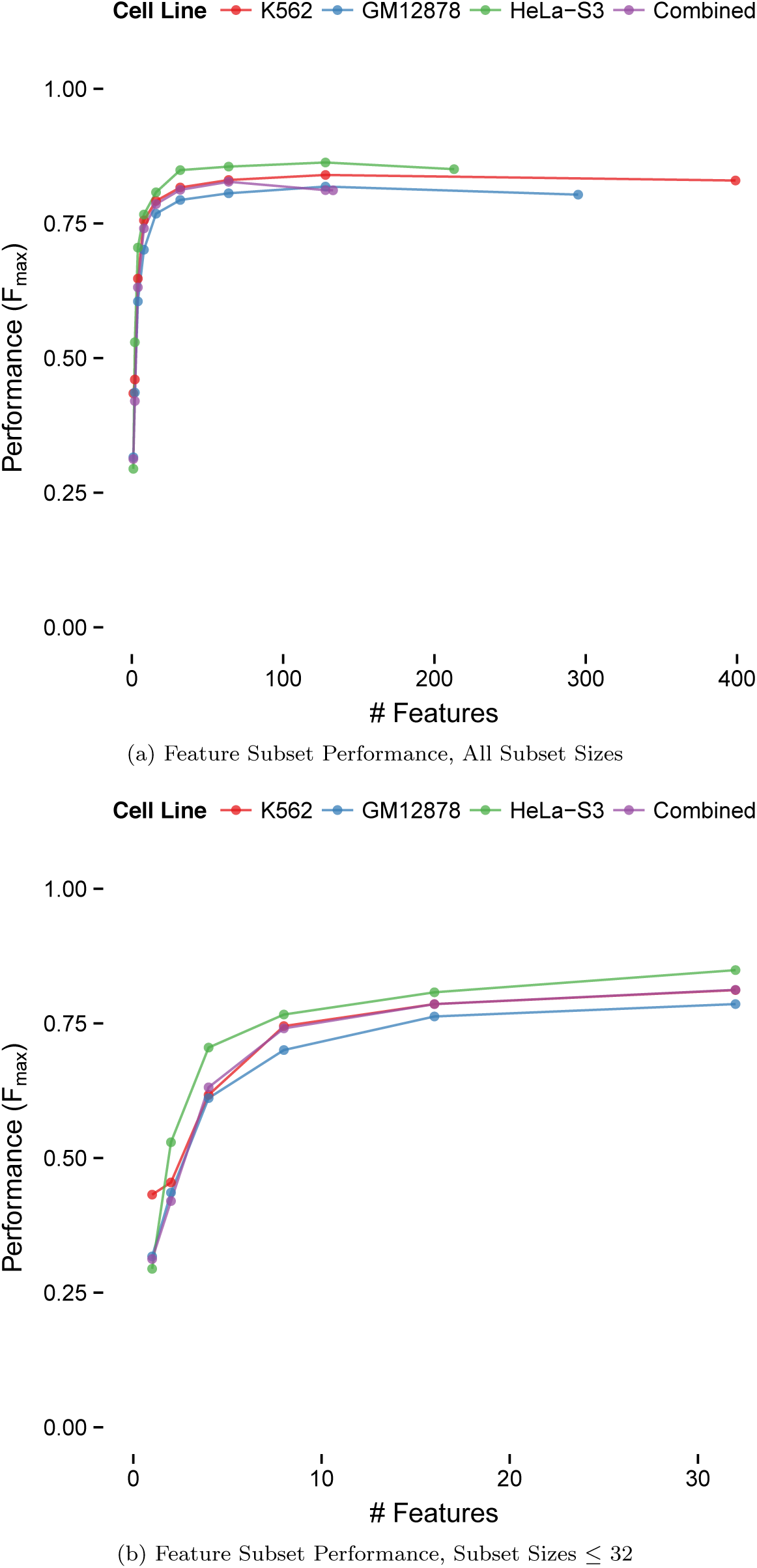
Performance as a function of the number of features. Performance of recursive feature elimination (Methods) that evaluates predictor subsets of size 1 up to the maximum per cell line and increasing by powers of 2 for computational efficiency. Near optimal performance was achieved using only ~16 predictors for lineage-specific models as well as the combined model, while lower but acceptable performance required only 8 predictors. Performance slowly degrades past a certain number of predictors due to the noise introduced by uninformative features. For extra detail, the second figure shows performance using the top 1, 2, 4, 8, 16, and 32 predictors.

## Discussion

Through precise chromatin looping, regulatory elements physically interact with promoters of their target genes over long genomic distances, while avoiding other nearby active and inactive promoters. How do they do this? We hypothesized that transcription factors, histones, and architectural proteins might combine to drive—or at least mark—distal regulatory interactions. If so, then we ought to be able to computationally model known interactions from functional genomics data, and the most important genomic datasets in the model might shed light on the mechanisms of gene regulation in three dimensions. To test this hypothesis, we built *TargetFinder*, a machine learning algorithm that finds an optimal combination of genomic features for predicting experimentally validated enhancer-promoter pairs. The resulting models of distal regulatory interactions achieved outstanding performance, with a balance of precision and recall across the K562, GM12878, and HeLa-S3 ENCODE cell lines ranging from 83-88%and an FDR up to fifteen times smaller than using the closest gene. Our findings were robust across multiple enhancer and promoter definitions, and were reproduced using multiple algorithms and programming languages.

### Which functional genomics experiments are most informative about chromatin interactions?

A unique feature of our approach is that we combine high resolution genome-wide Hi-C interaction data [33] with the vast functional genomics datasets provided by the ENCODE project for predicting distal enhancer targets. By integrating these diverse datasets and examining their relevance to enhancer-promoter interactions, we discovered the most predictive datasets and highlighted the complex interplay between regulatory proteins and DNA in the three-dimensional genome. All of the top ranking features were functional genomics experiments, rather than conserved synteny or similarity of TF and target gene annotations. We identified EBF1, FOXM1, NCOR1, PML, RFX5, SMARCC1, SRF, STAT1, TBP, and TCF7L2 as combinatorially predictive proteins whose role in distal enhancer-promoter interactions may be under-appreciated. Other predictive proteins that were independently predictive included CDS1, C/EBP*β*, GABPA, GATA2, HCFC1, JUN/JUNB/JUND, MEF2A, NFATC1, NFKB1, PHF8, REST, SAP30, SP2, SPI1, and THAP1. Many of these features interacted with the cohesin complex and ZNF143, which was recently shown to provide sequence specificity to cohesin-assisted chromatin looping [41]. Members of the cohesin complex (SMC3/RAD21), CTCF, and ZNF143 were also highly ranked and have greater potential to generalize across cell types than lineage-specific TFs. Such activators and repressors nonetheless boost performance in individual cell lines, particularly those related to AP-1 and PRC2 complexes. Well-known histone marks necessary for ChromHMM/Segway annotations of promoters and enhancers are also necessary, though we found activating marks H2AZ/H3K9ac and elongation marks H3K36me3/H3K79me2 were especially predictive.

### DNA between interacting enhancers and promoters carries a distinct genomic signature

The knowledge gained in this study depended critically on our decision to include genomic data from the window between each enhancer and promoter in the analyses. We discovered these window features dominated those encoding chromatin states at the promoter and enhancer themselves. The genomic signature of looping DNA had several components. First, interacting pairs tended not to have cohesin complex bound to the window, although it was prevalent near the enhancer and promoter. Long-range loops in the window between a candidate enhancer and promoter greatly reduced their interaction probability, suggesting pre-existing loops act as a kind of insulator between flanking elements. Secondly, DNA between interacting enhancers and promoters tended not to contain activating TFs and epigenetic marks of elongation and active transcription, all of which could indicate the presence of an alternative promoter target. On the other hand, windows did contain epigenetic marks associated with heterochromatin, polycomb-associated proteins, and co-factors of CTCF associated with its insulator function. Given this, our predictive features are more relevant to looping models of interaction than alternatives such as facilitated tracking [68]. Polycomb complexes appear to play several roles in distinguishing nearby targets. For example, PRC2-targeted CpG islands are enriched for REST and CUX1 binding motifs, both transcriptional repressors [69] with high predictive importance. In Drosophila, cohesin co-localizes with PRC1 at promoters and interacts to control gene silencing [70]. Given the conservation of PRC between flies and humans [71], this has implications for the interaction of cohesin and PRC for mammalian gene silencing and thus discrimination of target promoters. Also, distal enhancers may sometimes serve to clear PRC from CpG islands [72]. Finally, recent work shows that cohesin spatially clusters enhancers [73] and is consistent with our observation that the presence of active marks at alternate nearby enhancers often increase the likelihood of interaction. These are several of many possible explanations for the ability of window-based features to predict distal enhancer-promoter interactions with high precision and recall—explanations that may be refined by analysis of new functional genomics datasets.

### How does TargetFinder distinguish targets from non-target promoters in the same locus?

Careful examination of many enhancer-promoter pairs across cell lines suggests several broad rules influence *TargetFinder’s* score of an enhancer-promoter interaction: 1) do the enhancer and other nearby enhancers look active? 2) does the target transcript look like it is actively elongating? 3) is the target promoter cell type-specific? 4) do other promoters near the target have repressive marks or marks of paused polymerase? 5) is another pair interacting within the window? and 6) are there marks of chromatin remodelers or architectural proteins in the window, plus cohesion complex adjacent to the promoter and enhancer, that might facilitate looping interactions?

Figure 6 illustrates how these rules are combined to learn that an enhancer loops over the promoters of intervening genes (INTS6, WDFY2) to interact with the promoter of DHRS12 roughly 400 kilobases away. No single mark distinguishes the target. All active promoters have a repressive H4K20me1 mark and an activating (via Pol II elogination) H3K36me3 mark. Furthermore, PHF8 is present at every promoter in the region, while SP2 is present at none. Thus two highly predictive features do not separate targets from non-targets in this locus. Instead, the CTCF mark lacking cohesin complex marks suggests the WDFY2 promoter is not tethered to a distal enhancer via chromatin looping. However, DHRS12 has a cohesin complex mark (RAD21) at its promoter, and both RAD21 and SMC3 nearby. This interaction may also be defined by more complex interactions, including FOS and JUN binding on the looping chromatin, which is associated with changes in conformation [74, 75] that could possibly be relevant to the angle of the loop and not necessarily limited to their presence or absence at competing promoters. The situation appears even more complicated in loci with multiple active enhancers, including physically associated enhancers targeting the same promoter (Supplemental Figure S3). But *TargetFinder* can still predict enhancer-promoter pairs with high accuracy in such loci, indicating a degree of modularity in the genomic signature of interactions across loci, regardless of their architectures.

### Prospects for predicting regulatory interactions in many cell types

In addition to better understanding the mechanisms behind distal enhancer-promoter interactions within a specific cell line, we aimed to train *TargetFinder* on data-rich cell lines such as those provided by ENCODE, identify a minimal subset of easy to collect datasets needed for prediction, and make accurate predictions on new cell lines. Cross-cell line prediction is a difficult task as enhancers and promoters vary, functional genomics assays are noisy and may have different peak strengths due to numerous factors, and as few as 55% of interactions were shared between cell lines [33]. This condition is sometimes termed *covariate shift* and violates the assumptions of most machine learning methods. Despite these challenges, we discovered that accurate prediction requires only 8 ChIP-seq datasets, and nearly optimal prediction requires only ∼16. Importantly, many of these proteins are not routinely interrogated, and several frequently studied histones and TFs are redundant with or less predictive than proteins they interact with. Additionally, our analyses highlighted proteins that are predictive either in isolation or in combination with others. This impacts the probability that a dataset will generalize across cell types.

We therefore conclude that a researcher seeking to collect data for enhancer-promoter prediction in a new system might prioritize experiments that are in the top ~16 features (Figure 2) for the most similar well-characterized cell type or use features that score well across multiple cell lines. This will direct researchers towards predictive co-factors rather than multi-functional proteins that may be better known but less predictive. A *TargetFinder* trained on this reduced feature set from the well-characterized line(s) with validated interactions could then be applied to the new system by plugging in the values of the features for enhancer-promoter pairs in the new cell type, without the need for generating validated interactions. A subset of the resulting candidate interactions could then be tested using low-throughput assays to validate the predictions. In addition to reducing the burden on experimentalists, a model learned from fewer features is less likely to overfit and thus more likely to perform well on new cell lines. Our study demonstrates that this approach has the potential to be much more accurate than simply mapping enhancers to the closest promoter. Thus, *TargetFinder* is not only a tool for predicting the interactions of distal regulatory elements, but also a screening tool for estimating the relevance of unassayed DNA-binding proteins and epigenetic marks in disparate cell lines—and potentially disparate organisms.

## Materials and Methods

All code was implemented in Python using the `scikit-learn` machine learning library [76] and the pandas analytics library [77] in combination with `bedtools` [78]. Results were verified using a comparable pipeline implemented in R using the caret [79], `randomForest` [80], `gbm` [81], and glmnet [82] packages. Genome-wide data was obtained from the UCSC Genome Browser for the ENCODE Project [11] (http://genome.ucsc.edu/ENCODE/) for the K562 (tier-1), GM12878 (tier-1), and HeLa-S3 (tier-2) cell lines. GENCODE version 19 annotations and expression data were obtained directly from the ENCODE portal (https://www.encodeproject.org/data/annotations). Chromatin interaction data generated by Rao et. al [33] was obtained from the Gene Expression Omnibus (GEO). Training data was generated using the same methods and parameters for each cell line. When possible, separate features were generated for enhancer, promoter, and window regions defined as all base pairs between the proximal edges of the enhancer and the promoter.

### Promoter Identification

In each cell line, we identified actively transcribed protein coding genes with mean FPKM > 0.3 [84] and irreproducible discovery rate < 0.1 [85]. Corresponding promoters were regions labeled “TSS” (predicted promoter region including transcription start site) by the combined ChromHMM [86] and Segway [87] annotations available from the UCSC Genome Browser. This resulted in 9863, 10092, and 9303 active promoters for the above cell lines out of 20345 annotated protein coding genes. We also evaluated performance using GENCODE version 7 annotations and expression data, as well as promoter regions defined as a GENCODE TSS ± 2 kilobases (Supplementary).

### Enhancer Identification

Enhancers were segments labeled “E” (strong enhancer) by the combined ChromHMM [86] and Segway [87] annotations available from the UCSC Genome Browser. To focus our models on distal interactions, enhancers closer than 10 kilobases to the nearest promoter were discarded. This resulted in 44227, 51631, and 41734 active enhancers for the above cell lines. We also evaluated performance using clustered TF binding sites (Supplementary).

### Chromatin Interactions

Hi-C is an unbiased method for genome-wide identification of chromatin interactions [88]. Recent work [33] applied Hi-C with improved resolution to 9 cell types, 3 of which also had extensive ENCODE data. Hi-C interaction data obtained from GEO lists statistically significant interactions at 10% FDR, which we further filtered down to 1% FDR. Positive training samples were interactions with at least one active enhancer and at least one active promoter intersecting with forward and reverse Hi-C fragments. This resulted in 1100, 1368, and 855 interacting enhancer-promoter pairs for the above cell lines. We also evaluated performance using ENCODE 5C data (Supplementary).

Negatives were random pairs of active enhancers and promoters without a statistically significant Hi-C interaction. To select negatives matching the distribution of positive interaction distances, positives were assigned to one of 5 bins using quantile discretization of the distance between enhancer and promoter. For each bin, 5 negatives per positive were randomly selected for the training set. This number was chosen for computational efficiency, but cross-validated performance was similar using the complete set of negative enhancer-promoter pairs (*≈* 700-900k depending on the cell line).

### Features

Chromatin immunoprecipitation followed by sequencing (ChIP-seq) identifies where and how strongly TFs, architectural proteins, and modified histones bind along the genome. ChIP-seq assays for 141, 98, and 71 different proteins were performed genome-wide by the ENCODE consortium for the above cell lines. Peaks were called by ENCODE using a uniform pipeline and biological replicates where possible, then provided as BED files that specify peak locations and strengths. These were intersected with promoter, enhancer, and window regions. The average peak strength per region was used as a feature (computed as the sum of peak strengths divided by the region length in base pairs), resulting in 3 features per ChIP-seq dataset.

DNase I hypersensitive sites sequencing (DNase-seq) and Formaldehyde-Assisted Isolation of Regulatory Elements followed by sequencing (FAIRE-seq) are similar assays for identifying regulatory regions in the genome. These assays were converted to features using the above ChIP-seq methodology.

Reduced representation bisulphite sequencing (RRBS) identifies methylated DNA regions and was performed genome-wide by the ENCODE consortium for the above cell lines. These regions and their methylated base counts were intersected with our promoter, enhancer, and window regions. The percent of methylated bases within each region was used as a feature, resulting in 3 features.

Annotation-based features were derived from STRING [89], IMP [90], and GeneMANIA [91] by summing the interaction scores between the gene and all TFs predicted by CENTIPEDE to bind the enhancer [92]. Features for each tool were derived separately. A synteny-based feature was derived using the phylogenetic distance between enhancers and promoters covered by the same syntenic nets [93], summed over the 23 mammals in the UCSC Genome Browser 46-way multi-species alignment (http://hgdownload.cse.ucsc.edu/goldenPath/hg19/multiz46way/). Annotation and synteny features were excluded from our final trained version of *TargetFinder* due to their low predictive importance relative to their high computational cost.

### Machine Learning

Supervised ensemble learning algorithms [94] were used to predict enhancer-promoter interactions. Ensemble learning is a subfield of machine learning that trains multiple diverse models and combines their predictions to achieve performance greater than the best individual model. Supervised algorithms require labeled training data; enhancer-promoter pairs are labeled as *positive* if the regions interact according to Hi-C data (see above) and *negative* otherwise. We used both random forests [95] and gradient boosted trees [96] to ensure consistent results. The former constructs independent decision trees in parallel, while the latter iteratively constructs decision trees and places increasing emphasis on high-error samples.

For boosting, we used the `gbm R` package [81] and `GradientBoostingClassifier` in the `scikit-learn` Python package [76]. Nested cross-validation achieved optimal performance using 4096 iterations (trees), shrinkage (learning rate) 0.1, and interaction depth (maximum tree depth) 9. Similar performance was achieved with slightly more conservative parameters.

For random forests, we used the `randomForest` R package [80] and `RandomForestClassifier` in the `scikit-learn` Python package [76]. We used 1500 trees and left all other parameters at defaults. A much smaller forest achieved similar performance; the larger forest was used solely to stabilize estimates of feature importance.

We verified that the cross-validated performance and feature importances of boosting and random forests were similar, though differences are expected by design. In addition, we evaluated logistic regression (using `scikit-learn` [76]) and elastic nets tuned via nested cross-validation (using `caret` [79] and `glmnet` [82]). The resulting performance drop was substantial and emphasizes the importance of capturing non-additive feature interactions for predicting enhancer-promoter interactions. Baseline performance was estimated using random training labels.

For linear classifiers, features were first mean-centered and scaled to unit variance. For all classifiers, training samples within each CV fold were assigned weights inversely proportional to their class prevalence in order to compensate for severe class imbalance.

Feature importances given in the paper were estimated using only gradient boosting, for simplicity.

### Performance Evaluation

True positives (tp), false negatives (fn), false positives (fp), and true negatives (tn) are defined by the following contingency table comparing actual and predicted labels:

Our chosen classifiers generate scores representing confidence that a sample belongs to the positive class. To evaluate performance, scores above a threshold are given a positive label and otherwise are labeled negative. Raising this threshold results in fewer but more confident positive predictions. As a result, we used two metrics that summarize performance over all possible thresholds using the following base metrics:

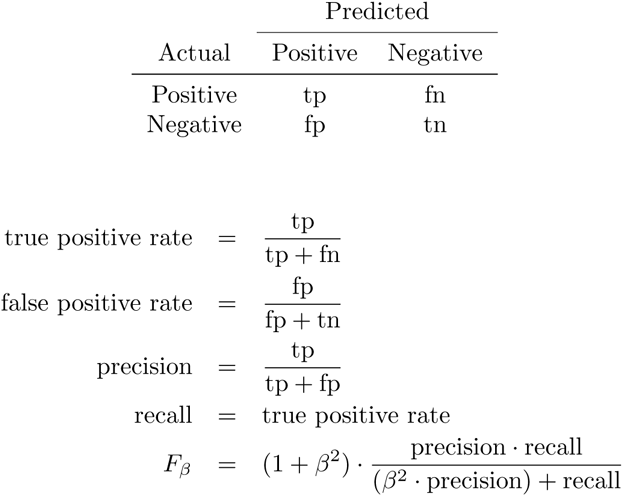

Area under the receiver operating characteristic curve (auROC or more commonly AUC) measures the area under the curve formed by the true positive and false positive rate of the classifier over all possible thresholds for the positive class. *F*_*β*_ is the weighted harmonic mean of precision and recall. We used the common *F*_1_ score (*F*_*β*_ where *β* = 1) to equally weight precision and recall. *F*_max_ is the maximum *F*_1_ score over all possible thresholds for the positive class.

AUC and *F*_max_ were estimated with 10-fold cross validation where data is split into 10 non-overlapping training and test sets. Samples were weighted with the inverse of their class counts to compensate for the imbalance between negative and positive samples. Classifiers were constructed for each training set and predictions were generated for the corresponding test set. Performance was evaluated independently for each test set and averaged to produce a single estimate per metric. For example, a 3-fold cross validation might result in an AUC of 0.7, 0.8, and 0.9 for folds 1, 2, and 3, resulting in an average AUC of 0.8.

### Ensemble Feature Selection

Recursive Feature Elimination (RFE) is an embedded multivariate feature selection technique [97] that has recently been adapted to random forests [98, 99]. The selection process is similar to parameter tuning where the best number of features is considered a parameter of the model, and requires nested cross-validation as detailed by Ambroise and McLachlan [100] to obtain unbiased performance estimates. Our analysis used `caret’s` implementation of RFE.

For each cross-validation fold, the ensemble is first trained using the complete feature set. Importances are then estimated by permuting the values of each feature over all out-of-bag samples and measuring the average loss in accuracy. Out-of-bag samples are those excluded at each tree during training as a result of resampling with replacement. The performance of feature subsets is then evaluated using inner cross-validation. The performance of the best subset from inner cross-validation is then re-estimated using only the outer test fold to avoid bias. The smallest subset size having average performance within 1.5% of the best performance across all folds was selected as optimal. To reduce computation time, we evaluated subset sizes from 1 up to the maximum number of features counting by powers of 2.

RFE performance was estimated using random forests, but not boosting, due to limitations in `caret`.

### Interaction Networks

A network of feature interactions was created using the top 10 predictive datasets per cell line. Nodes in the network were connected with an orange edge if they had Pearson correlation above 0.3 at enhancer, promoter, or window regions. Purple edges connected nodes with known protein interactions according to BioGrid 3.3.122 [101]. The the most central node (part of the most shortest paths between all nodes) were shown in bold. Central nodes often correspond to datasets with large rank changes in univariate versus multivariate performance. A limited number of nodes were shown to conserve space.

## Acknowledgments

This project was supported by the Bench to Bassinet Program of the NHLBI (U01HL098179), NIH/NHLBI (HL089707), the San Simeon Fund, and the Gladstone Institutes.

## Supplemental Material

### Alternate promoter and enhancer definitions

Before transitioning to annotation-based enhancers, we defined candidate enhancers as transcription factor binding sites (TFBS) identified by CENTIPEDE [92], lifted these over from the hg18 to hg19 assembly, and clustered them [102] using the DBSCAN algorithm [103] with `eps = 300` and `min_samples = 1`. Finally, we intersected the resulting TFBS clusters with p300, H3K27ac, and H3K4me1 ChIP-seq peaks from the same cell line and retained all clusters that overlapped at least one of these ChIP-seq marks. Clusters closer than 10kb to the nearest promoter were discarded. This approach had comparable performance for 5C-assayed interactions but was outperformed by annotation-defined enhancers for Hi-C-assayed interactions.

### Alternate interaction data

Before transitioning to Hi-C-assayed interactions, we used chromosome conformation capture carbon copy (5C) data from ENCODE that also identifies physically interacting segments of the genome [31]. Enhancers were intersected with forward 5C fragments and promoters with reverse 5C fragments. Following the ENCODE standard for interaction significance, enhancer-promoter pairs with fragments found to interact across both 5C biological replicates were given positive labels in our training data. To select negatives matching the distribution of interaction distances, positives were first assigned a bin number using quantile discretization of the distance between enhancers and promoters. For each positive distance bin, 200 negatives were generated by randomly selecting non-interacting enhancer-promoter pairs within the ENCODE pilot regions. The number of negatives per bin was limited by the number of active promoters covered by reverse 5C fragments. Due to the limited number of positives, we transitioned to Hi-C data when it became available with sufficient resolution.

### Supplemental Tables

**Table S1:**
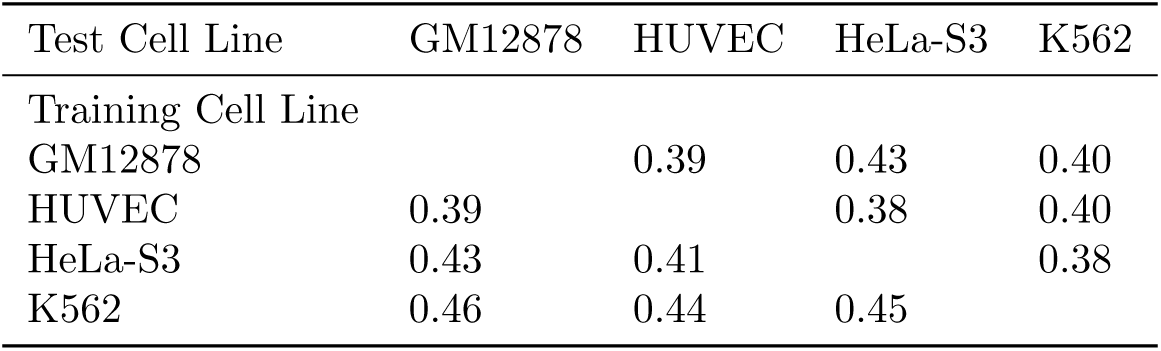
*TargetFinder* performance (*F*_max_) when trained on one cell line and tested against another. The top 16 features in the training cell line that were also present in the test cell line were used. An additional cell line not present in other evaluations, HUVEC, was used to test performance on a cell line where few (11) datasets were available.

| Test Cell Line | GM12878 | HUVEC | HeLa-S3 | K562 |
| --- | --- | --- | --- | --- |
| Training Cell Line |  |  |  |  |
| GM12878 |  | 0.39 | 0.43 | 0.40 |
| HUVEC | 0.39 |  | 0.38 | 0.40 |
| HeLa-S3 | 0.43 | 0.41 |  | 0.38 |
| K562 | 0.46 | 0.44 | 0.45 |  |

**Table S2:**
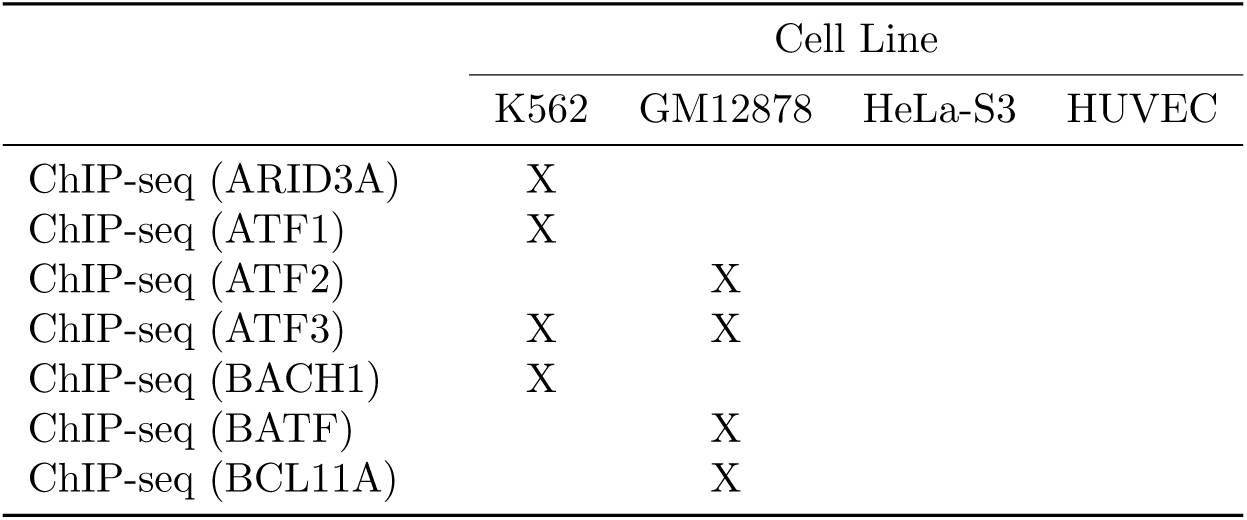
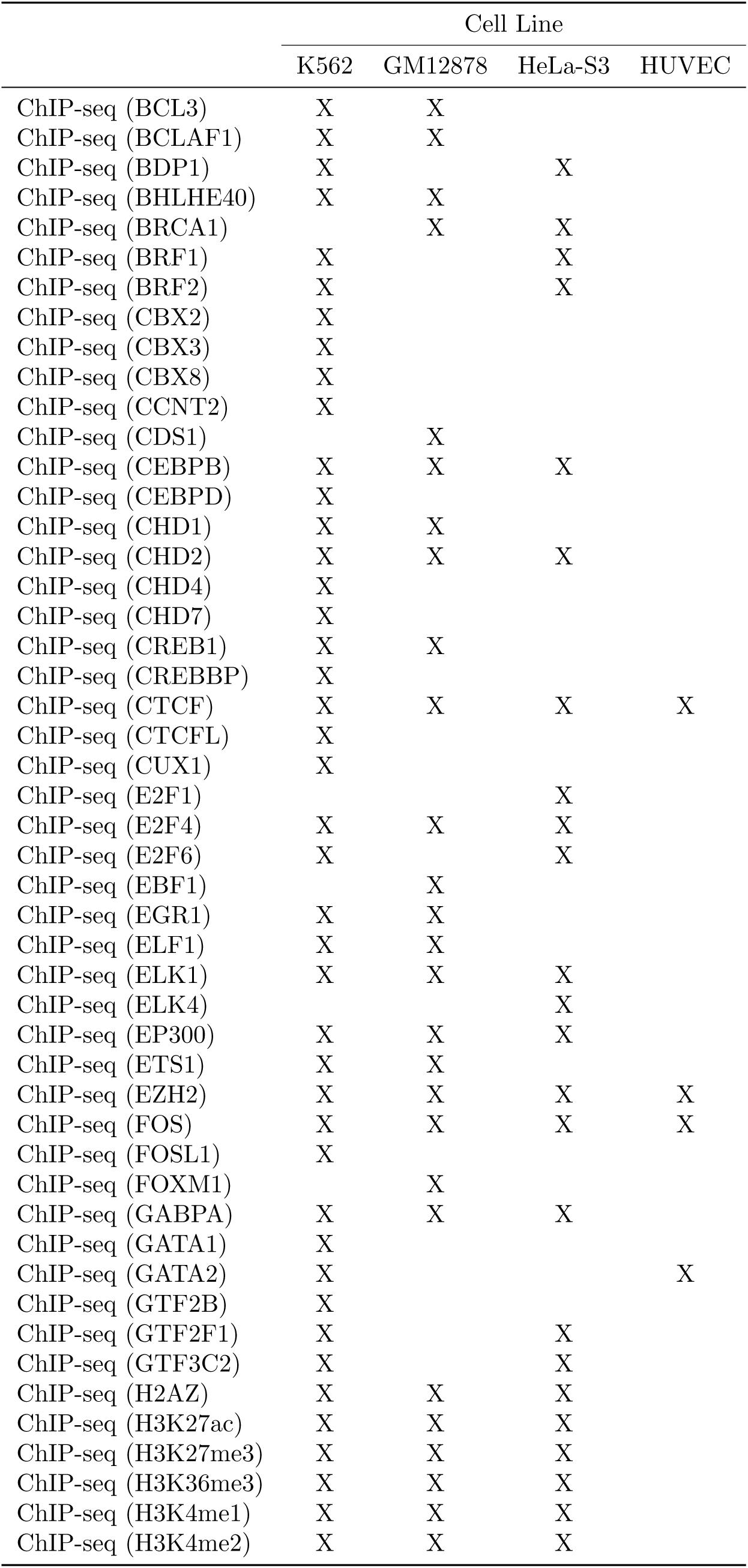
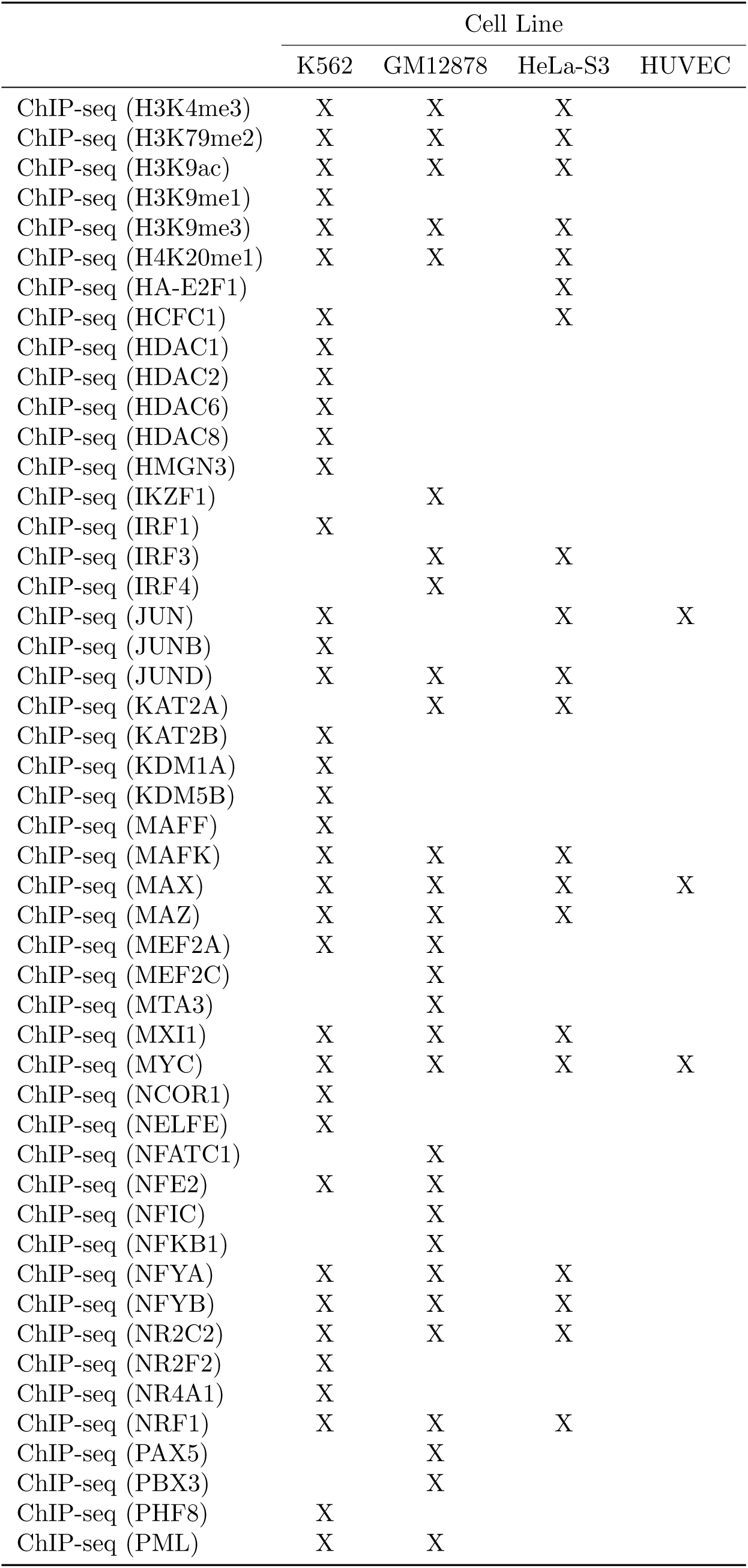
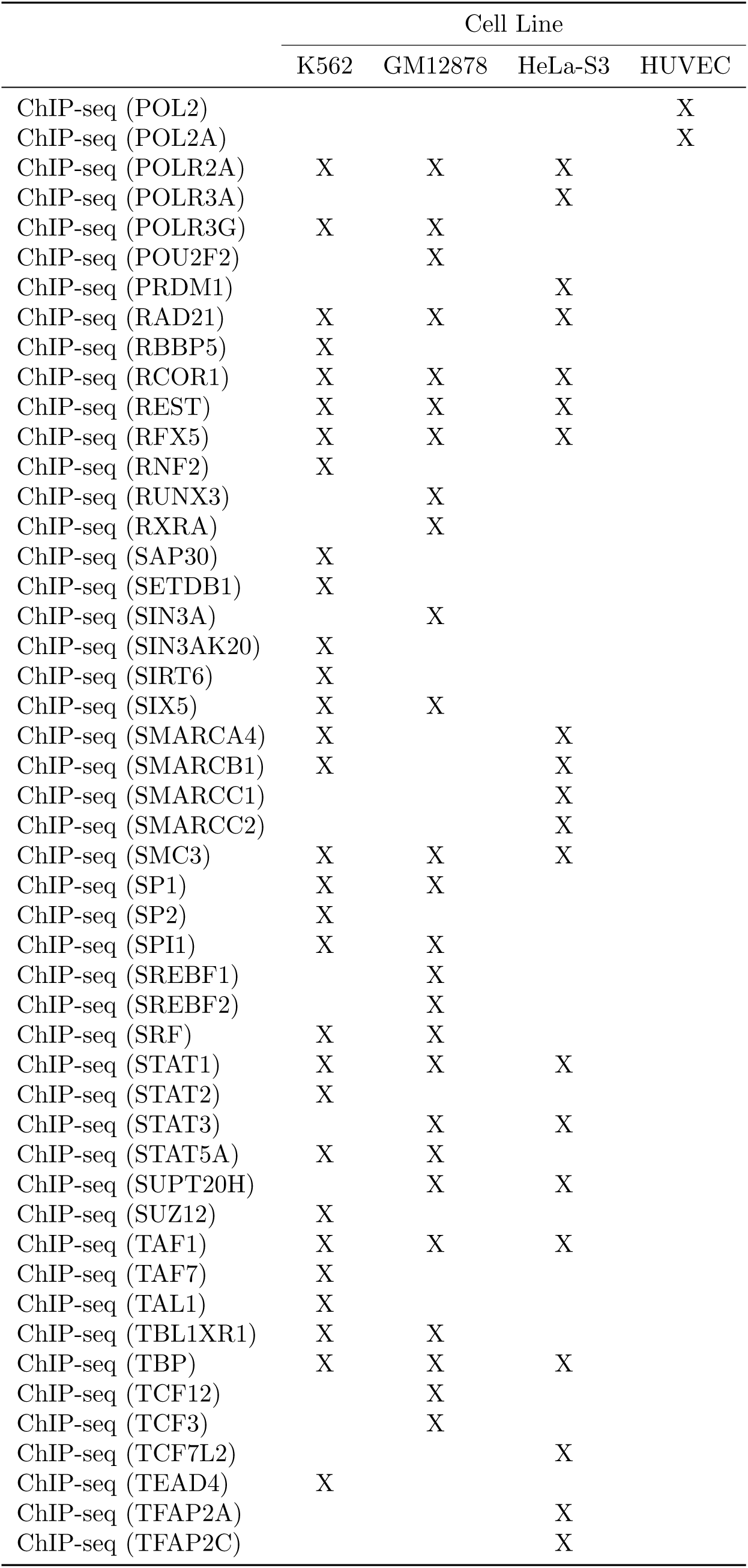
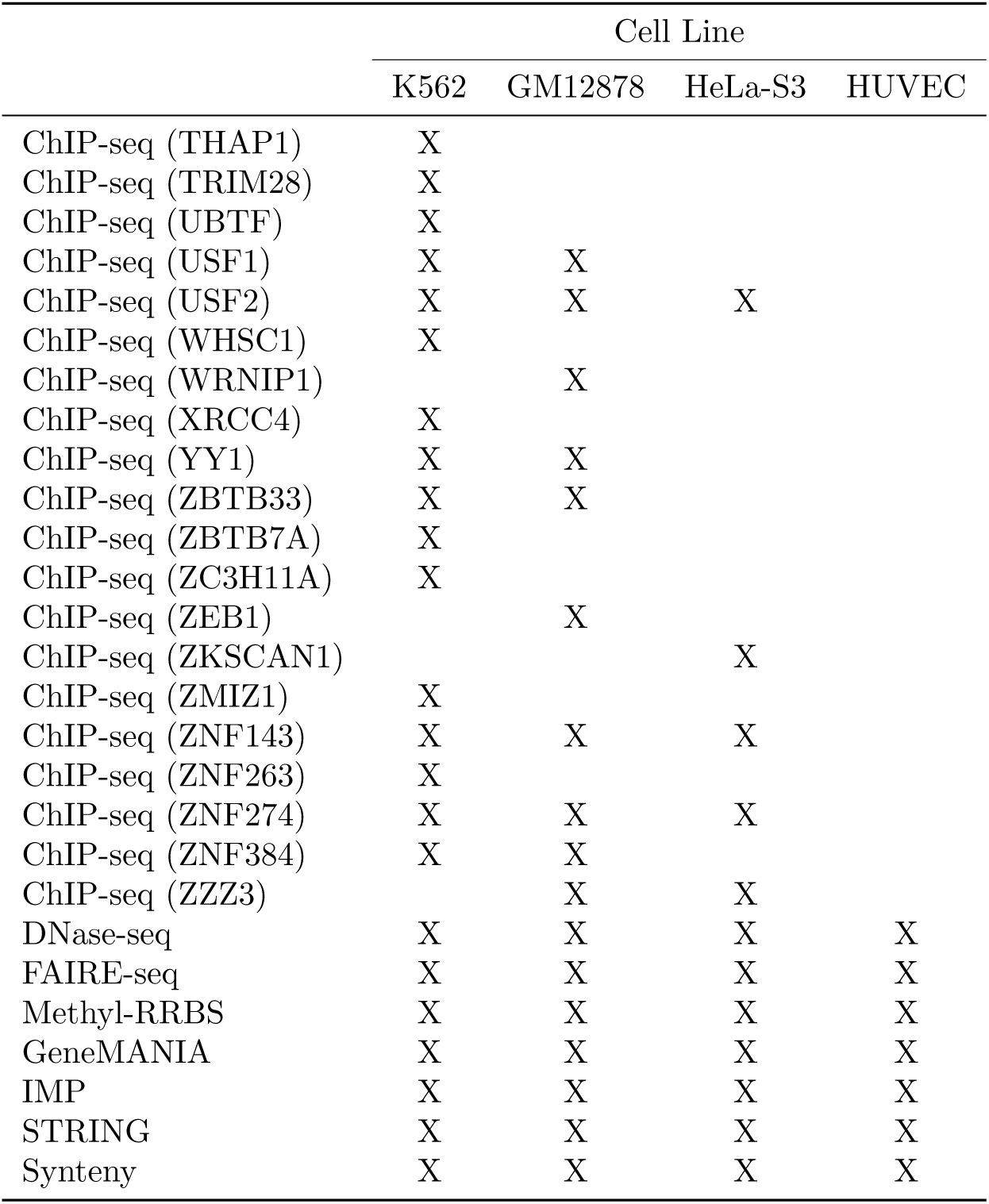
Datasets available for each cell line, driven primarily by availability from ENCODE.

**Table S3:**
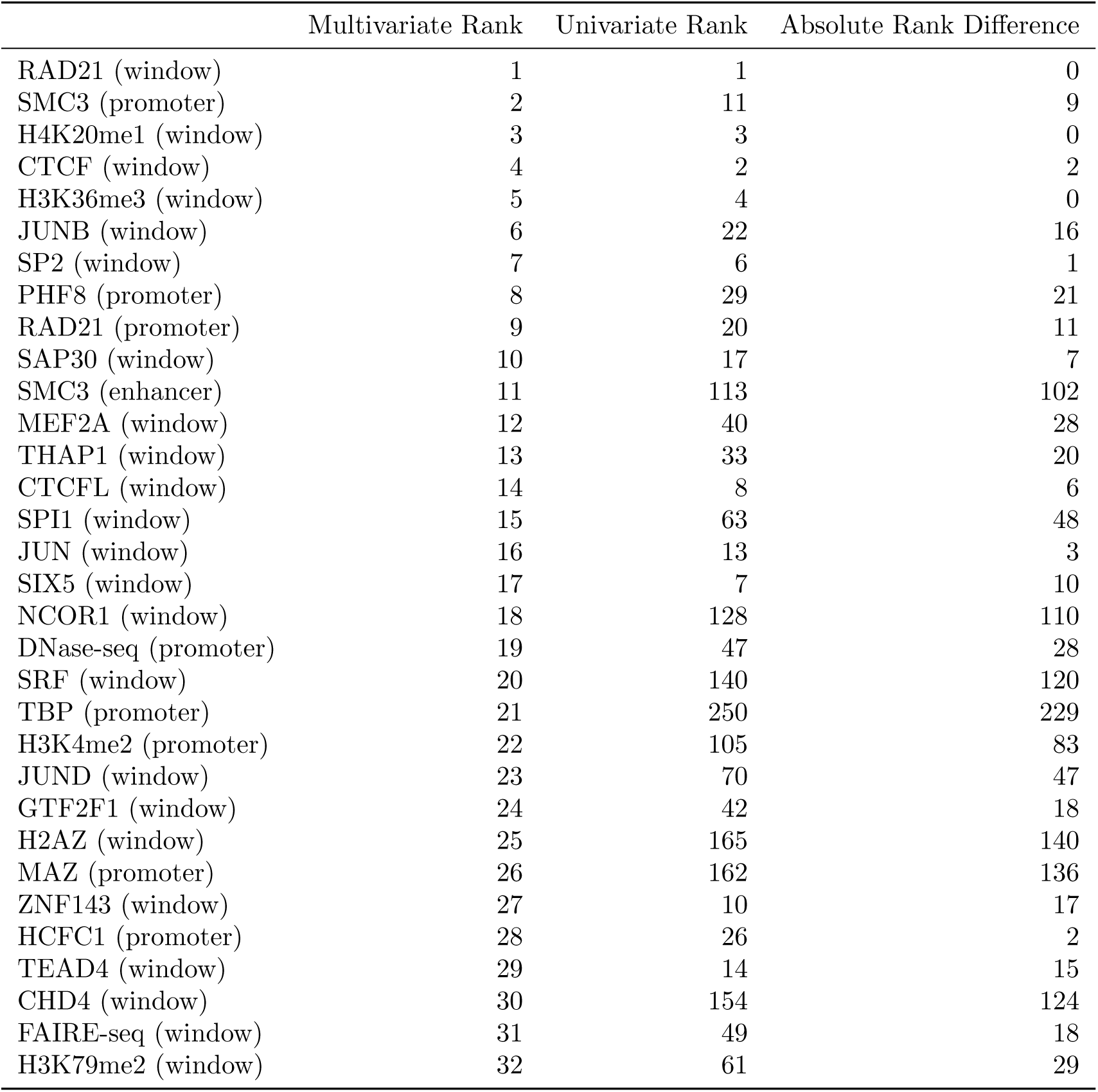
Univariate and multivariate feature ranks for the top 32 predictors in K562. Large differences between multivariate and univariate ranks indicate features that are not predictive on their own but become highly predictive in combination with other features. Such outliers may identify novel biological interactions or resolve ambiguities caused by noisy assays.

**Table S4:** Univariate and multivariate feature ranks for the top 32 predictors in GM12878. Large differences between multivariate and univariate ranks indicate features that are not predictive on their own but become highly predictive in combination with other features. Such outliers may identify novel biological interactions or resolve ambiguities caused by noisy assays.

|  | Multivariate Rank | Univariate Rank | Absolute Rank Difference |
| --- | --- | --- | --- |
| ZNF143 (window) | 1 | 1 | 0 |
| CTCF (window) | 2 | 2 | 0 |
| SMC3 (promoter) | 3 | 8 | 5 |
| RAD21 (window) | 4 | 3 | 1 |
| H3K9ac (enhancer) | 5 | 29 | 24 |
| H3K4me3 (promoter) | 6 | 19 | 13 |
| H2AZ (promoter) | 7 | 44 | 37 |
| FOXM1 (enhancer) | 8 | 82 | 74 |
| RAD21 (promoter) | 9 | 6 | 3 |
| H3K27ac (enhancer) | 10 | 46 | 36 |
| H3K9ac (promoter) | 11 | 57 | 46 |
| H3K9me3 (window) | 12 | 27 | 15 |
| H4K20me1 (window) | 13 | 53 | 40 |
| CDS1 (promoter) | 14 | 13 | 1 |
| NFKB1 (window) | 15 | 42 | 27 |
| RFX5 (window) | 16 | 37 | 21 |
| H3K4me3 (window) | 17 | 126 | 109 |
| H3K79me2 (window) | 18 | 60 | 42 |
| ATF2 (promoter) | 19 | 12 | 7 |
| DNase-seq (promoter) | 20 | 62 | 42 |
| BATF (window) | 21 | 28 | 7 |
| NFATC1 (window) | 22 | 69 | 47 |
| H3K4me1 (window) | 23 | 92 | 69 |
| H3K9ac (window) | 24 | 221 | 197 |
| NRF1 (window) | 25 | 5 | 20 |
| H3K36me3 (window) | 26 | 59 | 33 |
| H3K4me2 (promoter) | 27 | 39 | 12 |
| H3K27me3 (window) | 28 | 16 | 12 |
| EBF1 (window) | 29 | 158 | 129 |
| H3K79me2 (promoter) | 30 | 98 | 68 |
| H2AZ (enhancer) | 31 | 111 | 80 |
| REST (window) | 32 | 55 | 23 |

**Table S5:**
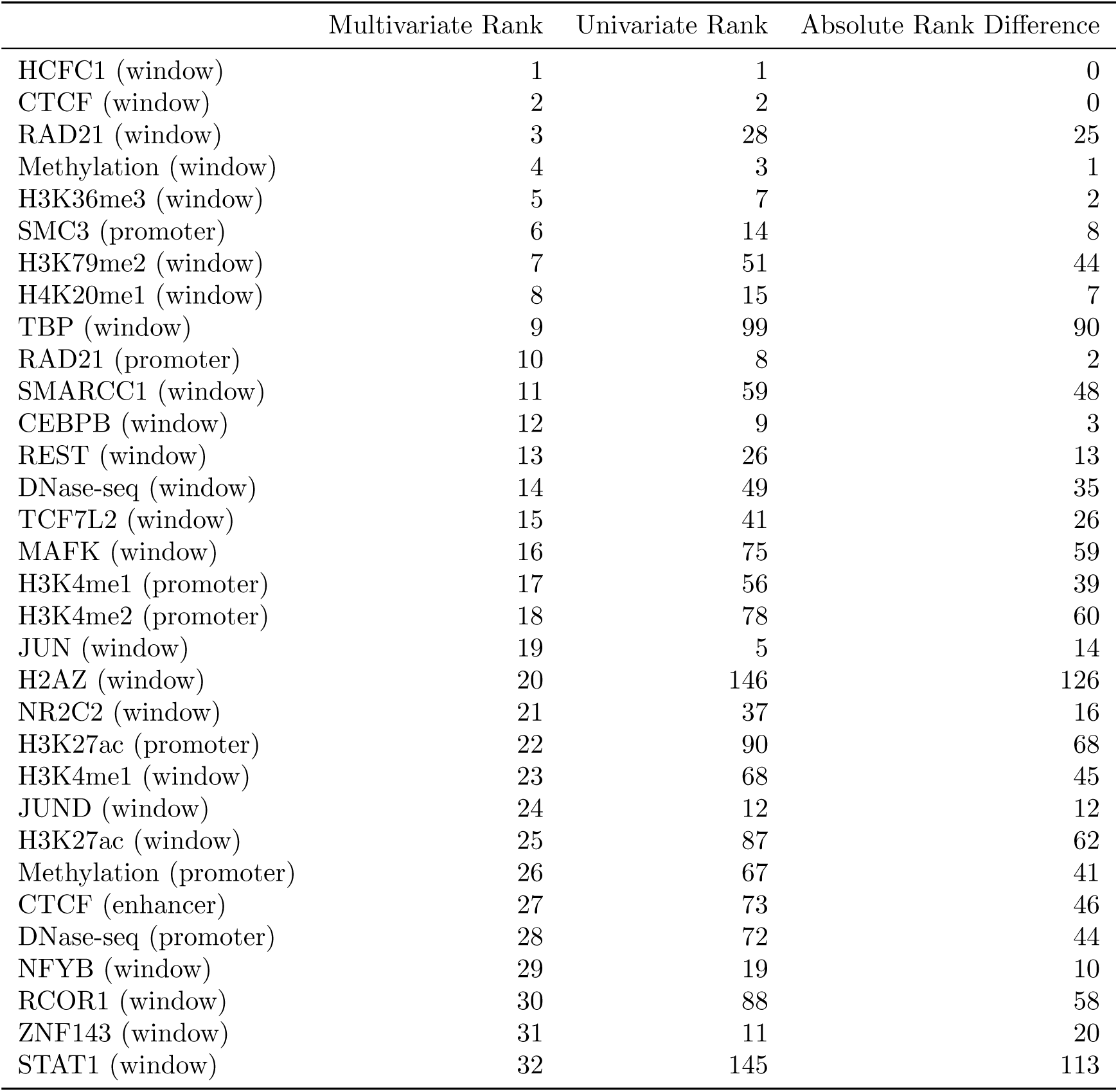
Univariate and multivariate feature ranks for the top 32 predictors in HeLa-S3. Large differences between multivariate and univariate ranks indicate features that are not predictive on their own but become highly predictive in combination with other features. Such outliers may identify novel biological interactions or resolve ambiguities caused by noisy assays.

|  | Multivariate Rank | Univariate Rank | Absolute Rank Difference |
| --- | --- | --- | --- |
| HCFC1 (window) | 1 | 1 | 0 |
| CTCF (window) | 2 | 2 | 0 |
| RAD21 (window) | 3 | 28 | 25 |
| Methylation (window) | 4 | 3 | 1 |
| H3K36me3 (window) | 5 | 7 | 2 |
| SMC3 (promoter) | 6 | 14 | 8 |
| H3K79me2 (window) | 7 | 51 | 44 |
| H4K20me1 (window) | 8 | 15 | 7 |
| TBP (window) | 9 | 99 | 90 |
| RAD21 (promoter) | 10 | 8 | 2 |
| SMARCC1 (window) | 11 | 59 | 48 |
| CEBPB (window) | 12 | 9 | 3 |
| REST (window) | 13 | 26 | 13 |
| DNase-seq (window) | 14 | 49 | 35 |
| TCF7L2 (window) | 15 | 41 | 26 |
| MAFK (window) | 16 | 75 | 59 |
| H3K4me1 (promoter) | 17 | 56 | 39 |
| H3K4me2 (promoter) | 18 | 78 | 60 |
| JUN (window) | 19 | 5 | 14 |
| H2AZ (window) | 20 | 146 | 126 |
| NR2C2 (window) | 21 | 37 | 16 |
| H3K27ac (promoter) | 22 | 90 | 68 |
| H3K4me1 (window) | 23 | 68 | 45 |
| JUND (window) | 24 | 12 | 12 |
| H3K27ac (window) | 25 | 87 | 62 |
| Methylation (promoter) | 26 | 67 | 41 |
| CTCF (enhancer) | 27 | 73 | 46 |
| DNase-seq (promoter) | 28 | 72 | 44 |
| NFYB (window) | 29 | 19 | 10 |
| RCOR1 (window) | 30 | 88 | 58 |
| ZNF143 (window) | 31 | 11 | 20 |
| STAT1 (window) | 32 | 145 | 113 |

**Table S6:**
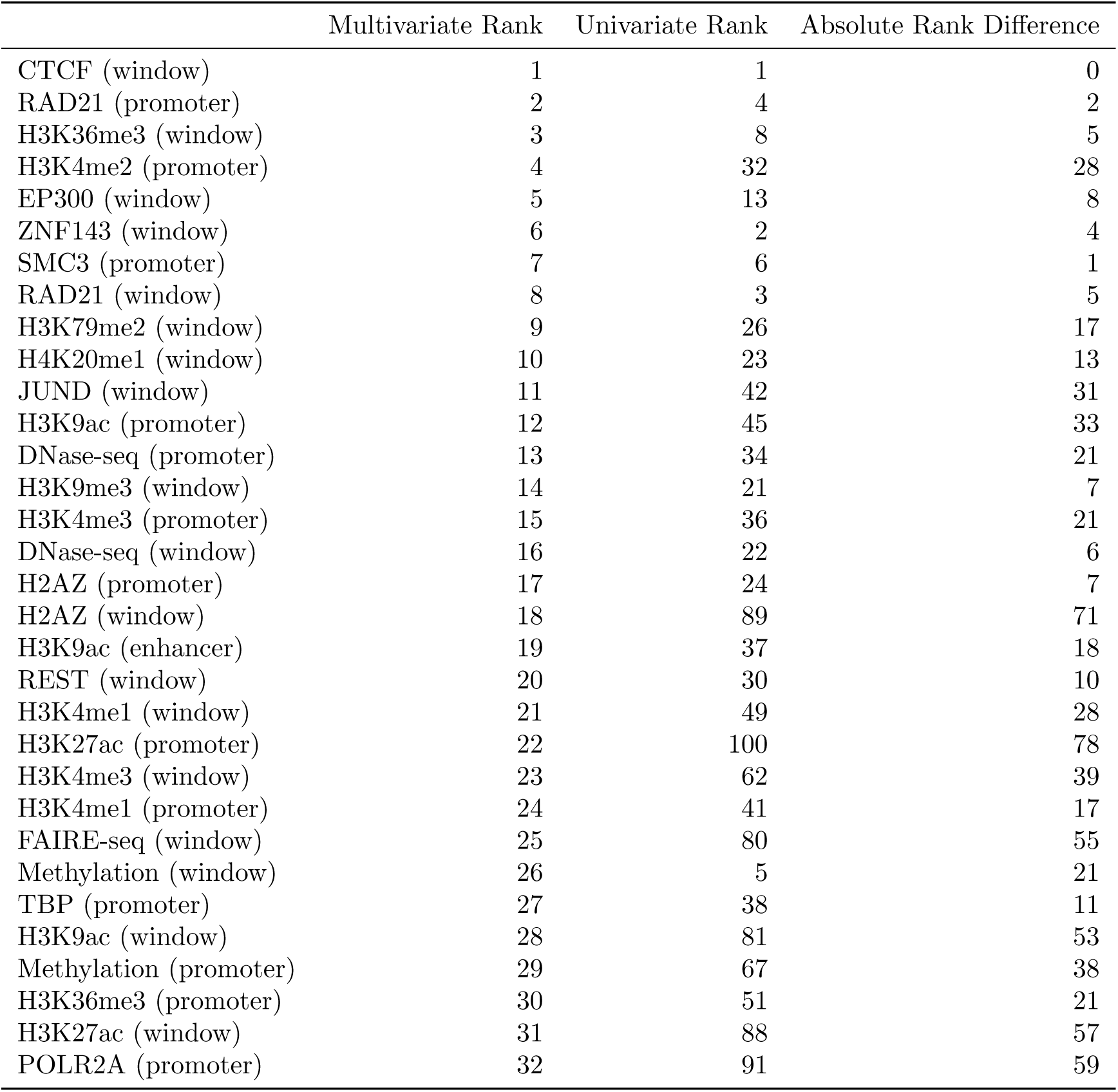
Univariate and multivariate feature ranks for the top 32 predictors across all cell lines. Large differences between multivariate and univariate ranks indicate features that are not predictive on their own but become highly predictive in combination with other features. Such outliers may identify novel biological interactions or resolve ambiguities caused by noisy assays.

|  | Multivariate Rank | Univariate Rank | Absolute Rank Difference |
| --- | --- | --- | --- |
| CTCF (window) | 1 | 1 | 0 |
| RAD21 (promoter) | 2 | 4 | 2 |
| H3K36me3 (window) | 3 | 8 | 5 |
| H3K4me2 (promoter) | 4 | 32 | 28 |
| EP300 (window) | 5 | 13 | 8 |
| ZNF143 (window) | 6 | 2 | 4 |
| SMC3 (promoter) | 7 | 6 | 1 |
| RAD21 (window) | 8 | 3 | 5 |
| H3K79me2 (window) | 9 | 26 | 17 |
| H4K20me1 (window) | 10 | 23 | 13 |
| JUND (window) | 11 | 42 | 31 |
| H3K9ac (promoter) | 12 | 45 | 33 |
| DNase-seq (promoter) | 13 | 34 | 21 |
| H3K9me3 (window) | 14 | 21 | 7 |
| H3K4me3 (promoter) | 15 | 36 | 21 |
| DNase-seq (window) | 16 | 22 | 6 |
| H2AZ (promoter) | 17 | 24 | 7 |
| H2AZ (window) | 18 | 89 | 71 |
| H3K9ac (enhancer) | 19 | 37 | 18 |
| REST (window) | 20 | 30 | 10 |
| H3K4me1 (window) | 21 | 49 | 28 |
| H3K27ac (promoter) | 22 | 100 | 78 |
| H3K4me3 (window) | 23 | 62 | 39 |
| H3K4me1 (promoter) | 24 | 41 | 17 |
| FAIRE-seq (window) | 25 | 80 | 55 |
| Methylation (window) | 26 | 5 | 21 |
| TBP (promoter) | 27 | 38 | 11 |
| H3K9ac (window) | 28 | 81 | 53 |
| Methylation (promoter) | 29 | 67 | 38 |
| H3K36me3 (promoter) | 30 | 51 | 21 |
| H3K27ac (window) | 31 | 88 | 57 |
| POLR2A (promoter) | 32 | 91 | 59 |

### Supplemental Figures

**Figure S1:**
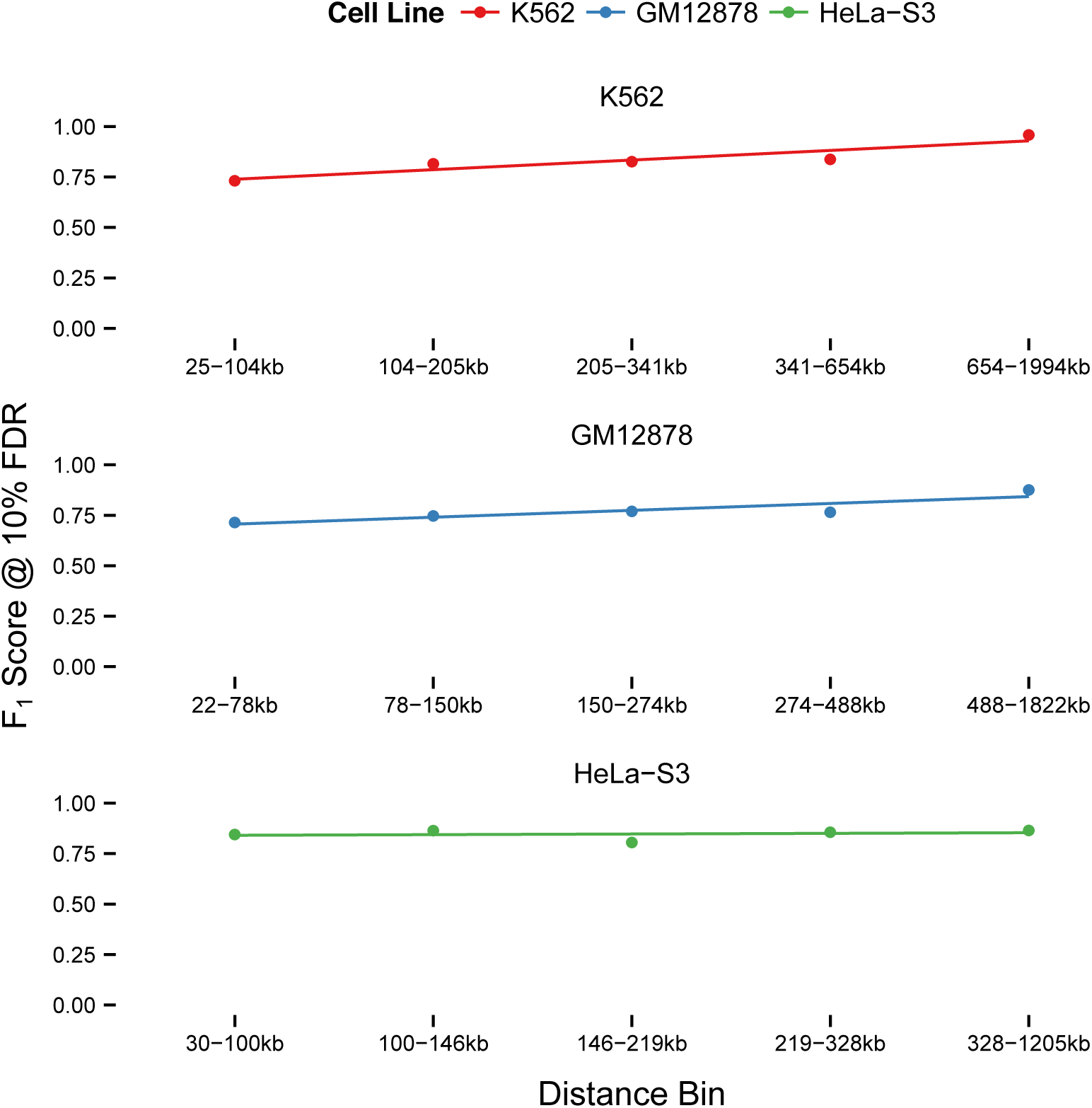
Performance as a function of distance between enhancer and promoter. Samples are first grouped into equal-count bins via distance quantiles, then predictions within each bin are evaluated at 10% FDR.

**Figure S2:**
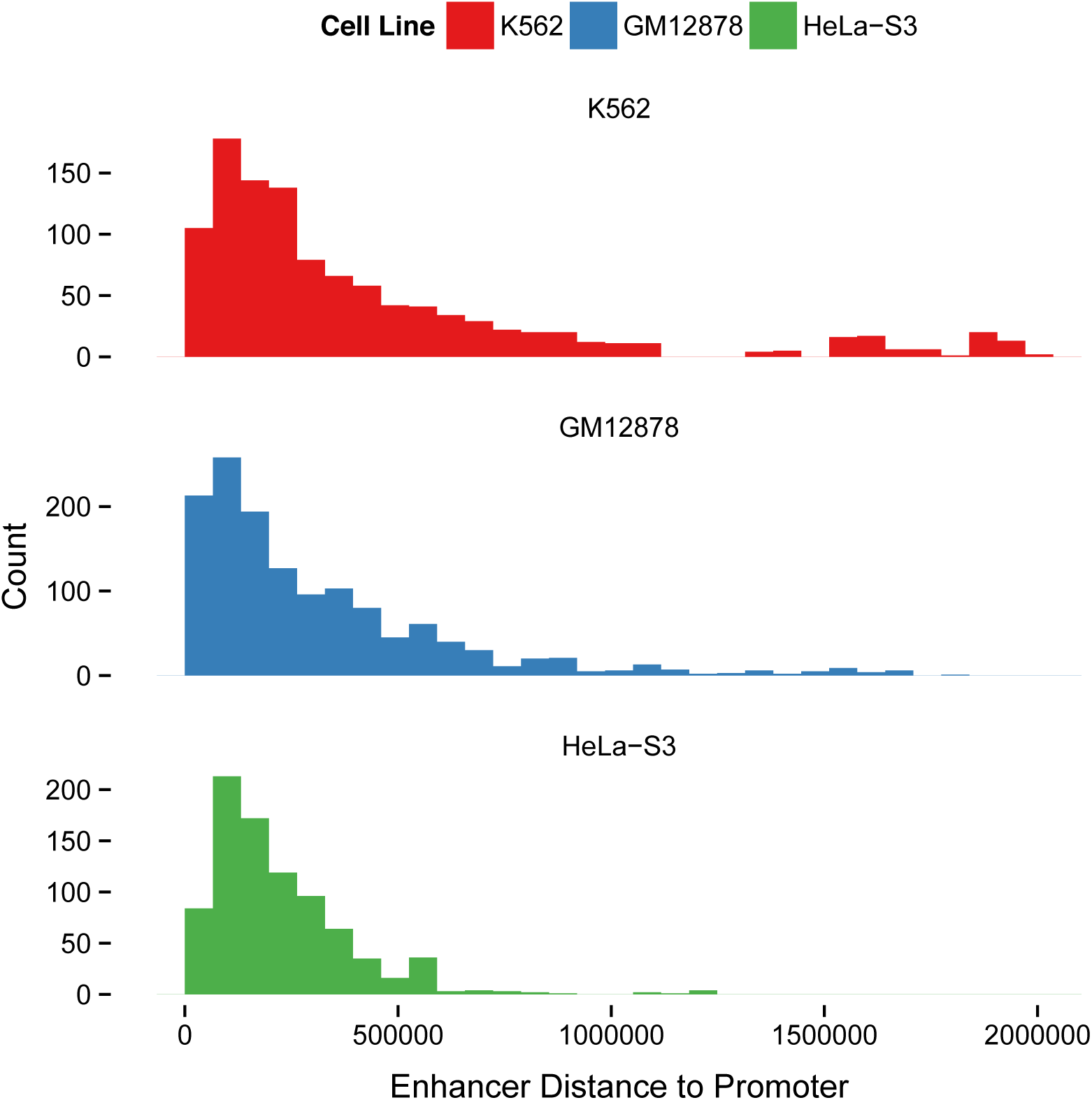
Distribution of genomic distance between enhancers and promoters in the training data for each cell line.

**Figure S3:**
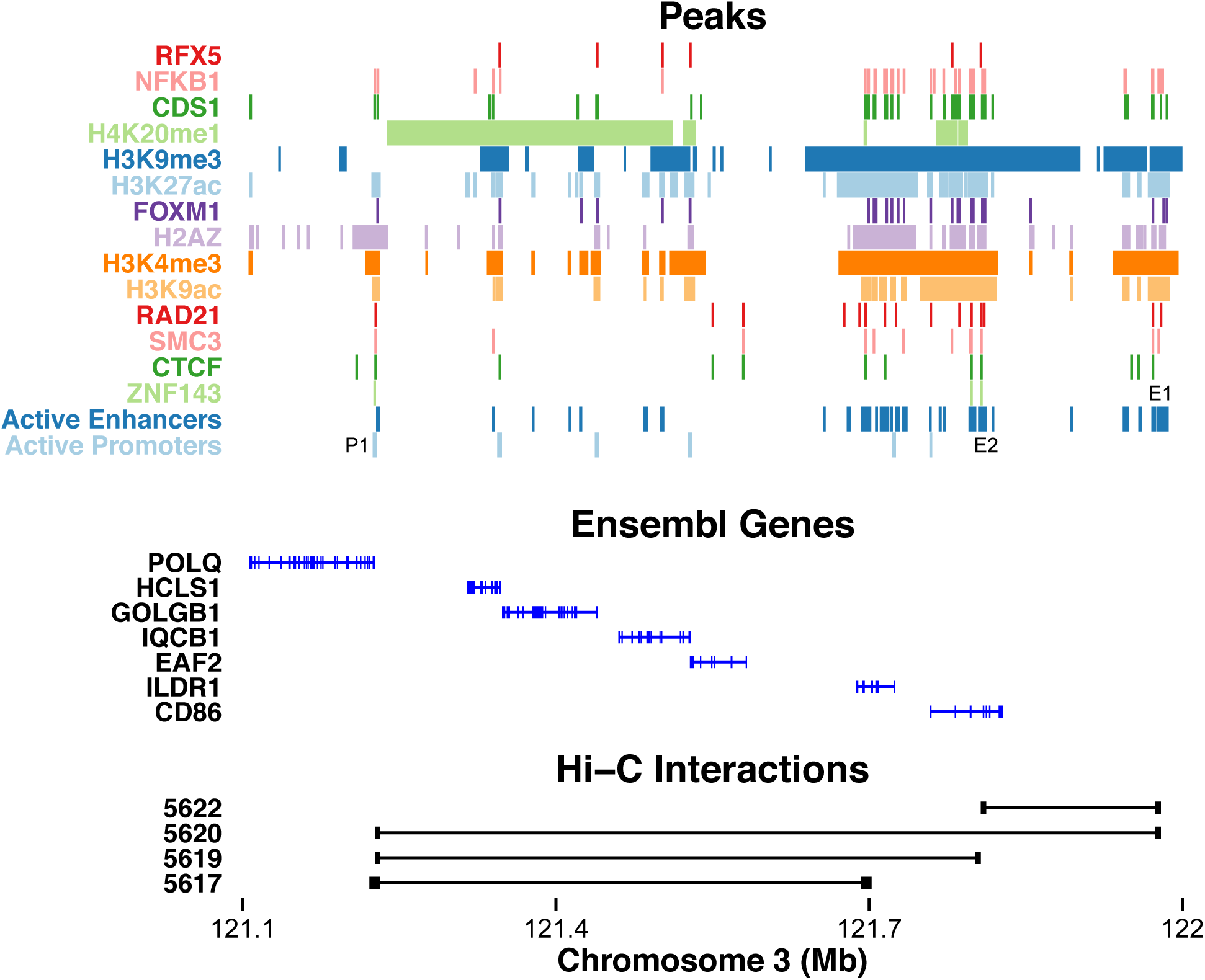
Significant peak values of the top 14 predictive datasets for an interacting promoter (P1) and enhancer (E1) in GM12878, separated by other active promoters and enhancers. In contrast to Figure 6, enhancer E1 interacts not only with P1 but also with E2, and P1 is the target of multiple enhancers. Active enhancers are segments marked “E” by combined ChromHMM/Segway annotations, and active promoters are segments marked “TSS” and expressed in GM12878 (determined by RNA-seq with expression threshold 0.3). Ensembl genes are also displayed, with introns denoted as thin lines and exons as squares. Left and right fragments of the Hi-C assay are also shown to visually confirm E1 interacts with P1 and other targets in the window. Note that P1 has all expected loop-associated marks including CTCF, cohesin, and ZNF143, as well as all other activation-associated marks. Also note spans of the repressive H4K20me1 and H3K9me3 marks that may rule out several alternate targets. As in the K562 example, the presence or absence of a single mark does not rule out a potential target and should instead be considered in combination with other marks.

